# Instantaneous visual genotyping and facile site-specific transgenesis via CRISPR-Cas9 and phiC31 integrase

**DOI:** 10.1101/2022.10.30.514454

**Authors:** Junyan Ma, Weiting Zhang, Zhengwang Sun, Saba Parvez, Randall T. Peterson, Jing-Ruey Joanna Yeh

## Abstract

The zebrafish *Danio rerio* has become a popular model in functional genomics and genetic disease studies. However, when a zebrafish mutant line must be propagated as heterozygotes due to homozygous lethality, using standard genotyping methods to identify a population of homozygous mutant embryos is time-consuming and sometimes impractical due to downstream applications such as large-scale chemical screens. Here, we introduce ‘TICIT’, <u>T</u>argeted <u>I</u>ntegration by <u>C</u>RISPR-Cas9 and <u>I</u>ntegrase <u>T</u>echnologies, which utilizes the site-specific DNA recombinase – phiC31 integrase – to insert fluorescent markers into CRISPR-Cas9-generated mutant alleles. It allows instantaneous determination of a zebrafish’s genotype simply by examining its color. This technique, which relies on first knocking in a 39-basepair phiC31 landing site via CRISPR-Cas9, enables researchers to insert large DNA fragments at the same genomic location repeatedly and with high precision and efficiency. We demonstrated that TICIT could also be used to create reporter fish driven by an endogenous promoter. Additionally, we created a landing site located in the *tyrosinase* gene that could support transgene expression in a broad spectrum of tissue and cell types, acting as a putative safe harbor locus. Hence, TICIT can yield predictable and reproducible transgene expression, facilitate diverse applications in zebrafish, and may be applicable to cells in culture and other model organisms.

## Introduction

The zebrafish is a genetically tractable vertebrate animal model that has been increasingly used for studying gene functions and genetic diseases in recent years (Demarest and Brooks-Kayal, 2018; Rissone and Burgess, 2018; Sakai et al., 2018). Zebrafish are fecund and economical, and they exhibit both physiological and pathological similarities to humans (Goldsmith and Jobin, 2012; Gut et al., 2017). The zebrafish genome shares high conservation with the human genome (Howe et al., 2013). It has been shown that 82% of disease-associated human genes have a zebrafish ortholog (Howe *et al*., 2013). Many biological phenomena have been shown to be highly conserved between zebrafish and humans at a genetic and mechanistic level (Cully, 2019; Demarest and Brooks-Kayal, 2018; Fazio et al., 2020; Torraca and Mostowy, 2018; Vacaru et al., 2014). As the scientific community seeks to assign functions to thousands of genetic variants being identified, the scale and throughput enabled by the zebrafish are likely to be invaluable. Moreover, zebrafish models permit chemical screens guided by successful reversal of disease-related phenotypes in a whole organism, which may substantially reduce discovery time and attribution rate during the development of therapeutics (Cui et al., 2011; Helenius and Yeh, 2012; MacRae and Peterson, 2015; Swinney and Anthony, 2011). Hence, developing genome engineering tools that can create zebrafish models more effectively and efficiently is expected to propel biomedical research that can later be translated into medicine (Cully, 2019; Swinney and Anthony, 2011).

While the zebrafish is highly amenable to genetic manipulations including Tol2-mediated transgenesis and CRISPR-Cas9-mediated mutations, the current methods are not without vexing shortcomings. Although transgenesis using the Tol2 transposon system is quite efficient, its outcome is often unpredictable and inconsistent between individual animals due to the variability in the number and location of integration event(s) (Abe et al., 2011). In contrast, in mice, Rosa26 has been extremely useful as a “safe harbor” genomic locus enabling faithful control of transgene expression by its own promoter (Rickert et al., 1997; Soriano, 1999). Single copy integration at the Rosa26 locus can be achieved by homologous recombination in embryonic stem cells or in mouse zygotes (Meyer et al., 2010; Rickert *et al*., 1997; Soriano, 1999). Nonetheless, to date, a defined safe harbor genomic locus has not been widely recognized in zebrafish. Various targeted knock-in methods via homologous recombination, microhomology-mediated end joining (MMEJ), or homology-independent mechanisms have been developed (Auer et al., 2013; DiNapoli et al., 2020; Hisano et al., 2015; Wierson et al., 2020). However, these techniques rely on endogenous DNA repair machinery, which may lead to variable efficiencies depending on the targeted loci and different repair outcomes for different integration events (Auer *et al*., 2013; DiNapoli *et al*., 2020; Hisano *et al*., 2015; Shin et al., 2014; Wierson *et al*., 2020). Currently, innovations and tools that allow zebrafish researchers to use a selected locus repeatedly for transgenesis are still lacking.

The DNA integrase of phiC31 bacteriophage is a site-specific recombinase that mediates DNA recombination between two heterotypic binding sequences named attB and attP, which are converted into two attB/attP hybrid sequences, termed attL and attR, after the recombination (Hillman and Calos, 2012). This reaction is irreversible, and thus phC31 recombinase can mediate stable integration (Hillman and Calos, 2012). Moreover, phiC31-mediated integration does not require any cellular auxiliary factors (Thorpe and Smith, 1998). It has been successfully used to insert large DNA constructs (10-100 kb) into the genomes of mammalian cells, fruit flies, *Xenopus*, and zebrafish (Belteki et al., 2003; Bischof et al., 2007; Li et al., 2012; Mosimann et al., 2013a; Roberts et al., 2014). Previously, Mosimann, *et al*. and Roberts, *et al*. generated phiC31 transgenesis recipient zebrafish lines via Tol2 and demonstrated that DNA vectors containing the attB sequence could be inserted into genomic attP sites, resulting in mean germline transmission efficiencies of 34% and 10% in the two studies (Mosimann et al., 2013b; Roberts *et al*., 2014). Encouraged by their results, we decided to explore additional applications using phiC31 integrase.

In this study, we combined CRISPR-Cas9 and phiC31 technologies and developed a workflow enabling facile single-copy, site-specific transgenesis at any user-specified targeted loci. Using this method called ‘Targeted Integration by CRISPR-Cas9 and Integrase Technologies’ or ‘TICIT’, researchers will be able to pre-select a suitable, well-characterized genomic location for inserting their transgenes. Given that multiple transgenic lines can be generated with insertions occurring at the same locus, expression differences due to different insertion sites can be avoided, and the expression levels of the transgenes among different transgenic lines are expected to be similar. Meanwhile, compared to other homology-directed approaches, this method eliminates the need for constructing homology arms into transgene vectors for each targeted locus. Instead, any vectors that contain a 70-basepair attB sequence can be used for integration. Researchers can also use TICIT to manipulate their genes of interest. Here, we applied this method to generate allele-tracking markers that can be seen even before the mutant phenotype manifests (Figure 1). This allows quick sorting of mutant embryos for any downstream applications. We also showed that TICIT can mediate in-frame integration enabling transgene expression controlled by an endogenous promoter (Figure 1). Finally, we characterized a safe harbor locus that may be useful for a variety of future studies (Figure 1).

**Figure 1.**
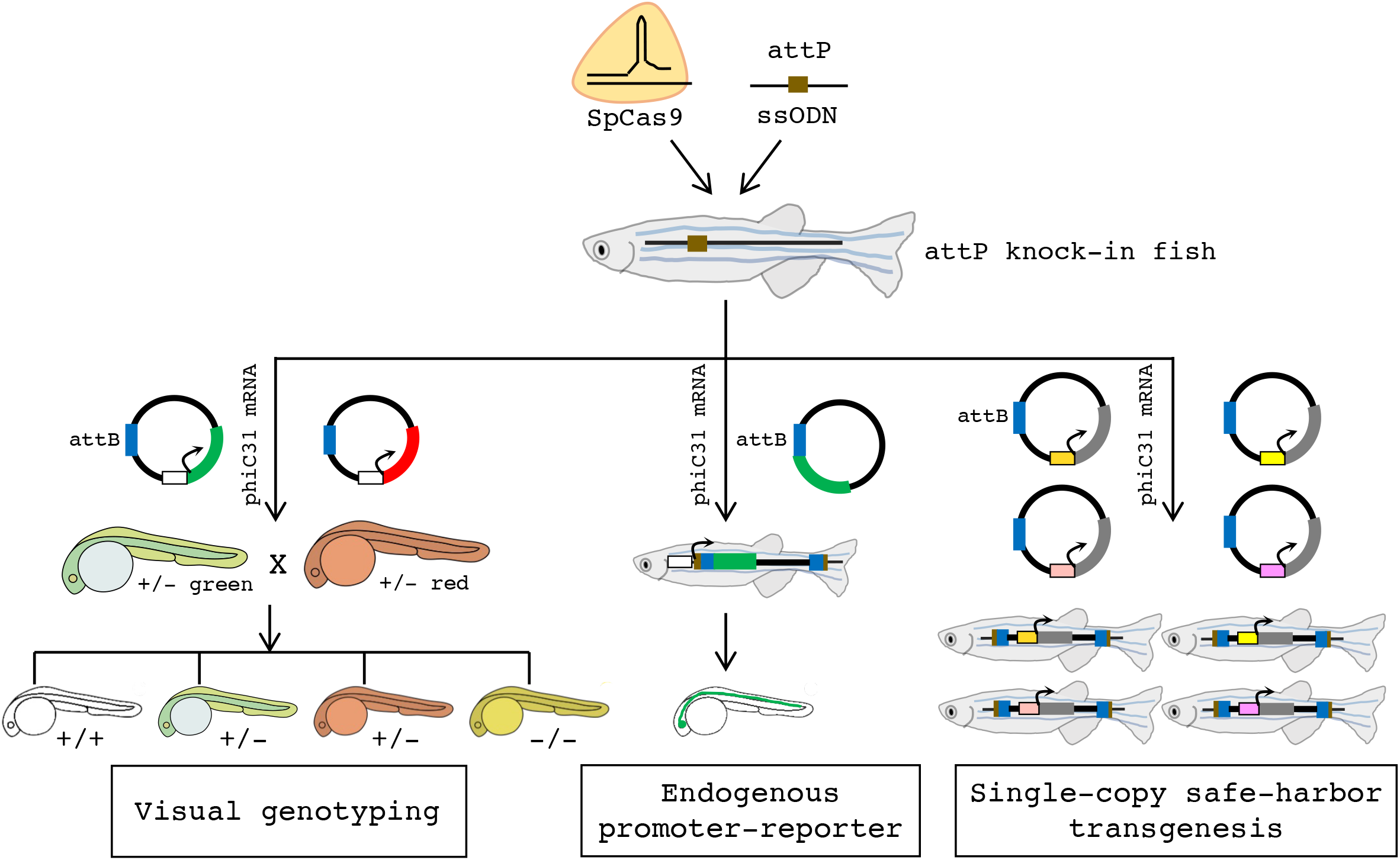
An overviewof the TICIT technology for visual genotyping, endogenous promoter-reporting, and single-copy, safe-harbor transgenesis. CRISPR-Cas9 from *Streptococcus pyogenes* (SpCas9) is employed together with a single-stranded DNA (ssODN) containing the phiC31 landing site to create attP knock-in fish lines. Next, single-copy site-specific transgene integration is achieved by co-delivering the phiC31 mRNA with any plasmids containing the phiC31 attachment sequence attB. For visual genotyping, two plasmids carrying fluorescent markers driven by a promoter of the user’s choice (shown as unshaded boxes) are used. This enables Cas9-mediated gene mutations to be identified by fluorescence even before mutant phenotypes manifest. In a heterozygous intercross, wild-type offspring are unmarked, heterozygous offspring are either green or red, and homozygous offspring are both green and red (shown as yellow). For endogenous promoter-reporting, a plasmid carrying a promoter-less fluorescent marker adjacent to the attB sequence is used. After phiC31-mediated integration, the promoter activity of the target gene can be tracked by the expression of the fluorescent marker. Further, fish lines carrying attP knock-in at pre-defined safe harbor loci may enable facile transgenesis devoid of positional and copy number artifacts.

## Materials and Methods

### Generation of gRNAs and Cas9 protein

All gRNAs used in this study were produced via in vitro transcription as previously described.(Gagnon et al., 2014) Target site and oligonucleotide sequences are listed in Supplemental Tables S1 and S2, respectively. Briefly, a gene-specific oligonucleotide was annealed to a constant oligonucleotide encoding the CRISPR-Cas9 scaffold and then a fill-in reaction was performed using T4 DNA polymerase (New England Biolabs). For the *tyr_1* and *tyr_2* sites, the 76-nucleotide (nt) SpCas9 (C9) scaffold was used. For gRNAs targeting *gfap* and *kcnh6a*, the 86-nt, enhanced SpCas9 (C9E) constant oligonucleotide was used.(Petri et al., 2022) The products were purified using Monarch^®^ PCR & DNA Cleanup Kit (New England Biolabs), which were then used as the template for in vitro transcription. In vitro transcription was performed using HiScribe™ T7 or SP6 High Yield RNA Synthesis Kit (New England Biolabs). RNA was subsequently purified using Monarch^®^ RNA Cleanup Kit (New England Biolabs).

For the generation of Cas9 protein, the plasmid pET-28b-Cas9-His (Addgene #47327) was transformed into Rosetta (DE3) competent cells (Novagen) following the manufacturer’s instruction. The production and purification of Cas9 protein were carried out as previously described.(Gagnon *et al*., 2014) Single-use aliquots were stored at −80°C.

### Plasmid construction

To generate the plasmid for *in vitro* transcription of the mouse codon-optimized phiC31 integrase (phiC31o) mRNA, a T7 promoter was added to the pPhiC31o plasmid (Addgene, #13794). Briefly, two oligonucleotides, EcoRI-T7 and T7-EcoRI (Supplemental Table S3), were hybridized with each other and ligated to EcoRI-linearized pPhiC31o. The construction (pT7_ PhiC31o) was transformed into NEB^®^ Turbo competent E. coli cells (New England Biolabs) and verified by Sanger sequencing after plasmid DNA extraction.

To generate pDestattB_ubi:mCherry, the mCherry coding sequence was PCR amplified from the plasmid pGFP-bait-MCS-NTR-mCherry using GoTaq^®^ DNA Polymerase (Promega) with primers listed in Supplementary Table S3. PCR products were purified using the Monarch^®^ PCR & DNA Cleanup Kit (New England Biolabs) and then digested by BspHI and MfeI-HF (New England Biolabs). Meanwhile, the plasmid pDestattB_ubi:EGFP (Addgene #68339) was digested by NcoI-HF and MfeI-HF (New England Biolabs) to generate the vector backbone. The digested mCherry and vector backbone fragments were purified using Zymoclean Gel DNA Recovery Kit (Zymo Research) and then ligated using the T4 DNA ligase (New England Biolabs). The construction was transformed into NEB^®^ Turbo competent E. coli cells (New England Biolabs) and verified by Sanger sequencing after plasmid DNA extraction.

To generate the pGEM-T-attB-P2A-EGFP plasmid, the EGFP coding sequence was PCR amplified from the plasmid pDestattB_ubi:EGFP using GoTaq^®^ DNA Polymerase (Promega) with attB-P2A-EGFP-F and pA-r primers (Supplementary Table S3). PCR product was purified and ligated to the pGEM^®^-T vector (Promega). The construction was transformed into NEB^®^ Turbo competent E. coli cells (New England Biolabs) and a partial truncation in the resulting clone was discovered by Sanger sequencing in the attB-P2A-EGFP-F primer region. Hence, we designed another primer attB-P2A-EGFP-F1 (Supplementary Table S3). The correct attB-P2A-EGFP sequence was PCR amplified from the plasmid with the truncated sequence using the attB-P2A-EGFP-F1 and pA-r primers. PCR product was purified and subcloned into pGEM^®^-T. The final construct pGEM-T-attB-P2A-EGFP was verified by Sanger sequencing.

### Oligonucleotide and mRNA synthesis

All oligonucleotides, including the ssODNs for attP knock-in, were ordered from Integrated DNA Technologies. The phiC31 integrase mRNA was prepared by *in vitro* transcription using *EcoRl*-linearized plasmid pCDNA3.1_phiC31 (Addgene #68310) or *HindIII-linearized* plasmid pT7_PhiC31o as the template and mMESSAGE mMACHINE™ T7 Transcription Kit (Invitrogen). The former vector contains a phiC31 coding sequence (phiC31)(Bischof *et al*.,2007), whereas the latter contains a mouse codon-optimized phiC31 coding sequence (phiC31o)(Raymond and Soriano, 2007). RNA was subsequently purified using Monarch^®^ RNA Cleanup Kit (New England Biolabs).

### Zebrafish microinjection

All zebrafish husbandry and experiments were approved by the Massachusetts General Hospital Subcommittee on Research Animal Care and performed in accordance with the guidelines of the Institutional Animal Care and Use Committee at the Massachusetts General Hospital.

Microinjections were performed using the one-cell stage of TuAB zebrafish embryos and approximately 2 nanoliters of injection solution per embryo. For attP knock-in experiments, the injection solution contained 480 ng/μl of Cas9 protein, 230 ng/μl of gRNA, and 0.5 - 1 μM of ssODN. To prepare the injection mix, Cas9 protein and gRNA were combined and put at room temperature for 5 minutes before the ssODN was added. We later used 288 ng/μl of Cas9 protein, 70 - 80 ng/μl of gRNA, and 0.5 uM of ssODN for the *gfap* and *kcnh6a* target sites to circumvent embryo death and deformity caused by high rates of *gfap* and *kcnh6a* mutations. For plasmid DNA integration experiments, the injection solution contained 12.5 ng/μl of the phiC31 or phiC31o mRNA and 12.5 ng/μl or 25 ng/μl of plasmid DNA (pDestattB_ubi:EGFP for the *tyr_1* and *tyr_2* targeted sites and pGEM-T-attB-P2A-EGFP for the *gfap* targeted site). We have used the mRNA of both phiC31 and phiC31o in these experiments and observed no consistent differences in their performance. For testing *gfap* and *kcnh6a* gRNA efficiencies, the injection solution contained 288 ng/μl of Cas9 protein and 80 ng/μl of gRNA. Injected embryos were incubated at 28.5°C after injection.

### Zebrafish genomic DNA extraction

Genomic DNA was extracted from fin clips of adult fish or embryos at 1 or 2 dpf. Zebrafish embryos that developed normally were lysed singly or as pools in lysis buffer (5 μl per embryo at 1 dpf, 8-10 μl per embryo at 2 dpf, and 30 μl per fin clip). The lysis buffer consisted of 10 mM Tris-HCl (pH 8.0), 2 mM EDTA (pH 8.0), 0.2% Triton X-100, and 0.5% Proteinase K.

Lysates were incubated at 50°C overnight with occasional mixing till they turned clear, which were then heated at 95°C for 10 mins to inactive Proteinase K. Genomic DNA was stored at 4°C.

### PCR-fluorescent fragment length (PCR-FFL) analysis and next-generation sequencing (NGS)

PCR-FFL analysis was employed to determine gRNA efficiencies or the sizes of the knock-in alleles.(Foley et al., 2009) To prepare the samples, two-step PCR reactions were performed. In the first step, gene-specific forward and reverse primers were used to amplify the targeted loci, and the forward primers contained an 18-bp M13 sequence (5’-TGTAAAACGACGGCCAGT) at the 5’ end. The PCR product was diluted 100-fold and used for the second PCR reaction using the 5’ 6-FAM-labelled M13 forward primer and a gene-specific reverse primer. PCR primer sequences are listed in Supplemental Table S3. The final products were analyzed at the Massachusetts General Hospital DNA Core.

For NGS, PCR amplicons (generally less than 280 bps) encompassing the targeted loci were generated using 1 μl of the zebrafish lysate with Phusion^®^ High-Fidelity Polymerase (New England Biolabs) and primers listed in Supplemental Table S3. PCR products were purified using the Monarch^®^ PCR & DNA Cleanup Kit (New England Biolabs) and submitted to the Massachusetts General Hospital DNA Core. Sequencing data were analyzed with CRISResso2 using the HDR mode (http://crispresso2.pinellolab.org/submission).

### Zebrafish genotyping, line generation, and founder screens

The sequences of all PCR primers for genotyping are listed in Supplemental Table S3. To generate attP knock-in lines, embryos microinjected with SpCas9, gRNA, and attP knock-in ssODN were raised to maturity and screened for founders. Potential founders (F_0_) were outcrossed to the wild-type fish, and their progeny (F_1_) were genotyped in pools (5 embryos per pool) via two-step nested PCR. The first PCR was to amplify the targeted loci, and the second PCR was to detect the attP insertion at the targeted loci (Supplemental Table S3). Alternatively, when F_1_ progeny were lysed individually, only the second step of PCR was needed to detect the attP insertion. Once an attP insertion was detected by PCR, the knock-in alleles were further verified by Sanger sequencing or next-generation sequencing, all using gene-specific primers to amplify the targeted loci (Supplemental Table S3). For sequence confirmation by Sanger sequencing, PCR products were subcloned using the pGEM-T vector, followed by colony PCR to identify the clones containing the desired allele. Subsequently, the plasmid DNA was extracted and submitted to the Massachusetts General Hospital DNA Core for sequencing. For genotyping of F_1_ and F_2_ adult zebrafish, fish were anesthetized briefly with tricaine, and a small fin biopsy was taken for DNA extraction as described above. Gene-specific primers were used to amplify the targeted loci (Supplemental Table S3), and the samples containing the knock-in alleles should yield PCR products corresponding to both the wild-type allele and the knock-in allele. For further verification, Sanger sequencing was employed.

To generate allele-tracking reporter lines, embryos from heterozygous *attP^tyr_1^* fish incrosses or outcrosses with the wild-type fish were injected with the phiC31 mRNA together with pDestattB_ubi:EGFP or pDestattB_ubi:mCherry DNA. The injected embryos were raised to adulthood and screened for founders. The progeny of potential founders was first screened for fluorescence. Fluorescent F_1_ embryos were lysed singly, and PCR was performed to detect DNA integration at the targeted locus (Supplemental Table S3). Sanger sequencing was used to confirm the sequences of attL and attR at the junctions of the integrated DNA. Subsequently, fluorescent progeny (F_1_) from the confirmed founders were raised to adulthood and genotyped by fin clipping and PCR as described above. We used two criteria to determine that a heterozygous *ubi:EGFP^tyr^* F_1_ fish did not carry a second integration outside of the targeted locus. First, when a heterozygous fish was outcrossed to a wild-type fish, it should produce approximately 50% of the fluorescent progeny. Second, all fluorescent F_2_ progeny should harbor the correct integration. F_1_ fish that fit these criteria were used to propagate all future generations.

To generate a target gene-specific reporter line, embryos from heterozygous *attP^gfap^* fish outcrossed to wild-type fish were injected with the phiC31 mRNA together with pGEM-T-attB-P2A-EGFP DNA. The injected embryos were raised to adulthood and screened for founders. The founder screen was performed as described for creating allele-tracking reporter lines.

### Confocal imaging of the gfap:gfp transgenic embryos

Heterozygous *P2A-EGFP^gfap^* embryos were obtained by mating heterozygous *P2A-EGFP^gfap^* fish with wild-type TuAB fish. EGFP-positive embryos were selected for imaging at one- and two-days post fertilization (dpf). Embryos were treated with 0.03 g/L phenylthiourea (PTU) to inhibit pigmentation, and manually dechorionated using tweezers. Right before imaging, embryos were anesthetized by 30 mg/L tricaine-S (Western Chemical) and mounted in 2% low melting agarose (LONZA) for dorsal or lateral view in 35 mm Petri dishes with a glass bottom. Imaging was performed on ZEISS LSM900 confocal microscope using a 10X objective. Z-stacks were collected with an interval of 3 μm. Images were stitched and computed as the sum of projections in ImageJ.

### Histology and cytology

Adult zebrafish were euthanized using approved protocols with tricaine and then fixed in 4% paraformaldehyde (PFA). Paraffin embedding, sectioning, hematoxylin & eosin (H&E) staining, and immunohistochemistry (IHC) for EGFP were performed using standard protocols. EGFP was detected using the JL-8 mouse monoclonal antibody (Clontech). For IHC, the primary antibody was diluted at 1:200, and the secondary antibody, an HRP-conjugated, donkey anti-rabbit antibody (Biovision), was used at a dilution of 1:1000.

For flow cytometry analysis of hematopoietic cells, *ubi:EGFP^tyr^* adult zebrafish were euthanized prior to kidney collection. The kidney was dissected and placed into ice-cold 0.9X phosphate-buffered saline (PBS) containing 5% fetal bovine serum. Whole kidney marrow (WKM) cells in single-cell suspension were generated by gently triturating and by passing through a 40-um filter. FACS was conducted on CytoFLEX (Beckman Coulter, NJ, USA) and data were analyzed with FlowJo software (Tree Star, OR, USA). Various hematopoietic cell populations were identified as previously reported (Traver et al., 2003).

## Results

### Generation of genomic attP landing sites using CRISPR-Cas9

To insert attP into targeted genomic loci, we used *Streptococcus pyogenes* Cas9 (SpCas9) to create double-strand DNA breaks and used single-stranded oligonucleotides (ssODNs) as the donor DNA for DNA repair. This technique has been used to create designer mutations in zebrafish (Boel et al., 2018; Jao et al., 2013; Petri *et al*., 2022). Meanwhile, using this method, we have previously shown that small precise edits can be introduced at an allele frequency up to ~10% in the injected embryos (Petri *et al*., 2022). We first targeted two SpCas9 cleavage sites in the *tyrosinase (tyr)* gene, named *tyr_1* and *tyr_2*. Both guide RNAs (gRNAs) can efficiently mutate *tyr* and elicit the albino phenotype (Jao *et al*., 2013; Moreno-Mateos et al., 2017). We designed the ssODNs to contain an attP flanked by two short homology arms adjacent to the SpCas9 cleavage sites (Figure 2A and Supplemental Table S4). For attP, a 39-basepair (bp) minimal sequence that showed full recombination activity in human cells and had little or no effect on transgene expression was used (Calos, 2006; Kirchmaier et al., 2013; Mosimann *et al*., 2013b). The attP sequence was inserted in different orientations in the ssODNs for these two loci so that both knock-in alleles would possess an in-frame stop codon in *tyr*. Further, the sequences for the homology arms were complementary to the non-target strand of SpCas9 and were 36 nucleotides (nts) long on the protospacer adjacent motif (PAM)-distal side and 91 nts on the PAM-proximal side. We and others have previously shown that this donor DNA configuration is effective in zebrafish and human cells (Petri *et al*., 2022; Prykhozhij et al., 2018a; Richardson et al., 2016). The ssODNs were chemically synthesized and two phosphorothioate linkages were added to both termini to enhance stability and knock-in efficiency (Prykhozhij et al., 2018b).

**Figure 2.**
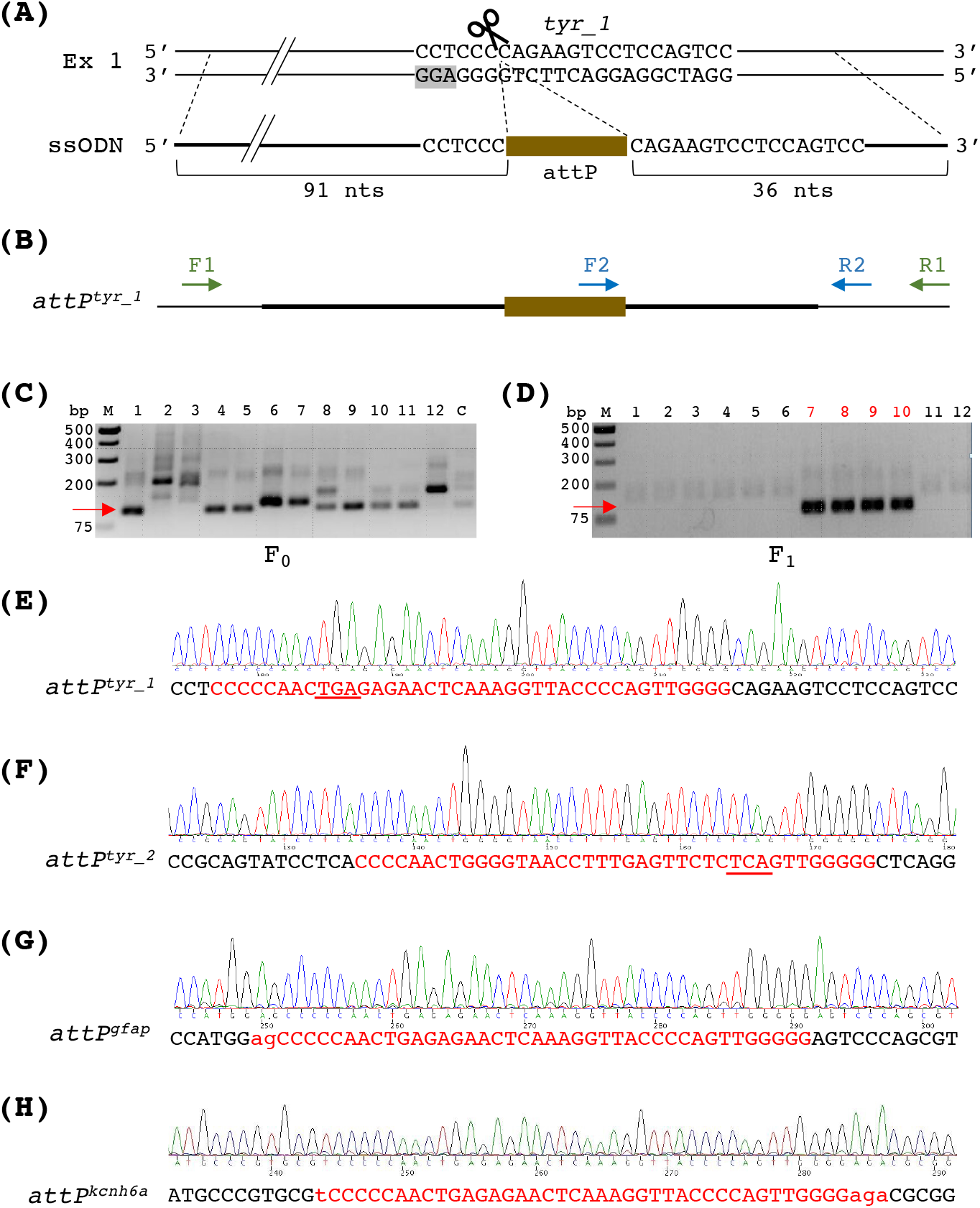
Generation of the attP knock-in fish lines. (A) Schematic of the SpCas9 target locus for *tyr_1* and its corresponding ssODN. The *tyr_1* site is in the first exon (Ex 1) of the *tyr* gene. Its target sequence is shown, and the spacer adjacent motif (PAM) is highlighted in gray. The cleavage site of SpCas9 is indicated by a scissor. The ssODN used for attP knock-in harbors attP encompassed by two homology arms. The sequences for the homology arms were complementary to the non-target strand of SpCas9 and were 36 nucleotides (nts) long on the PAM-distal side and 91 nts on the PAM-proximal side. Complete sequences of the ssODNs for attP knock-in at both *tyr_1* and *tyr_2* sites can be found in Supplemental Table S1. (B) Schematic of the attP knock-in at the *tyr_1* locus (designated as the *attP^tyr^_^1^* allele) and the primers for PCR analysis. The thick black line indicates the overlapping sequence with the ssODN. To detect the attP insertion, nested PCR was performed using *tyr*-specific F1 and R1 primers, followed by F2 and R2 primers that recognize attP and the *tyr* gene, respectively. (C) Representative agarose gel analysis of the nested PCR from single embryos microinjected with SpCas9 protein, the *tyr_1* gRNA, and the ssODN. The expected product size is 110 base pairs (bps) as indicated by a red arrow. (D) Representative agarose gel analysis of the nested PCR from the *attP^tyr_1^* founder screen. Twelve pools of embryos (5 embryos per pool) from each potential founder were analyzed. The 110-bp band (red arrow) indicated the presence of the *attP^tyr_1^* allele. The pools containing the attP knock-in alleles are labeled by red numbers on top. (E-H) Sequences of the attP knock-in at the *tyr_1* (E)*, tyr_2* (F)*, gfap* (G), *and kcnh6a* (H) targeted loci in the F_1_ fish. Endogenous gene sequences are in black letters. Insertions are in red letters. The attP sequences are in uppercase. For *attP^tyr_1^* and *attP^tyr_2^*, in-frame stop codons are underlined. For *attP^gfap^* and *attP^kcnh6a^*, the lowercase letters are sequences added to keep the insertions in-frame for unperturbed translation of the endogenous proteins.

We performed microinjection of 1-cell stage zebrafish embryos with SpCas9 and gRNA ribonucleoprotein (RNP) complexes and the ssODN for each sgRNA. Subsequently, successful knock-in events were identified in two ways. First, we conducted two-step nested PCR to detect the attP integration in the *tyr* gene. As illustrated in Figure 2B for the *tyr_1* locus, a pair of *tyr*-specific primers (denoted as F1 and R1) located outside the region of the donor DNA was used for the first round of nested PCR to amplify the targeted locus. We used another *tyr*-specific primer (denoted as R2) and an attP-specific primer (denoted as F2) for the second-round PCR to amplify the attP knock-in alleles. This yielded PCR products of not only the expected but also incorrect lengths, suggesting that some knock-in alleles may contain additional insertions or deletions (Figure 2C). Second, PCR-positive samples were chosen and their amplification products generated using a set of *tyr*-specific primers were subjected to next-generation sequencing. The results showed that this method had successfully inserted the entire attP site into both targeted loci (Table 1).

**Table 1.** Next-generation sequencing (NGS) demonstrates precise attP knock-in at the targeted loci. NGS was used to verify attP knock-in sequences in a pool of embryos previously injected with SpCas9 protein, gRNA, and ssODN. Total read counts and the percentages of different types of alleles are shown. ‘Unedited alleles’ are reads containing unmodified wild-type sequences. ‘Precise attP knock-in alleles’ are reads containing the expected knock–in sequences without any mutations. ‘Byproducts’ are reads containing imprecise attP knock-in sequences or indel mutations without attP knock-in sequences. For *tyr_1*, genomic DNA from five embryos that tested positive for attP knock-in by PCR were pooled together. For *tyr_2*, genomic DNA from two embryos that tested positive for attP knock-in by PCR were pooled together. For *kcnh6a* and *gfap*, genomic DNA from ten randomly selected embryos were pooled together. Genomic DNA

| Target site | Total read counts | Unedited alleles | Precise attP knock-in alleles | Byproducts |
| --- | --- | --- | --- | --- |
| *tyr_1* | 56146 | 0.19% | 9.11% | 90.70% |
| *tyr_2* | 97410 | 2.25% | 5.14% | 92.61% |
| *gfap* | 27950 | 7.18% | 0.80% | 92.02% |
| *kcnh6a* | 54464 | 65.21% | 0.05% | 34.74% |

We raised the injected embryos to adulthood, then outcrossed them to wild-type fish and screened for founders. The same nested PCR strategy was employed to identify the embryos carrying the attP knock-in alleles (Figure 2D). Further, the attP knock-in sequences in the F_1_ embryos were verified via Sanger sequencing (Figure 2E-F). For *tyr_1*, we identified one founder from 16 F_0_ fish screened. For *tyr_2*, we identified one founder from 3 F_0_ fish screened. Thus, founder frequencies were 6.3% and 33.3% for *tyr_1* and *tyr_2*, respectively. Meanwhile, germline mosaicism of the identified founders was 5.5% - 11.9% (Table 2). When F_1_ fish reached adulthood, we identified heterozygous fish carrying the attP knock-in alleles (hereafter named the *attP^tyr_1^* and *attP^tyr_2^* alleles) by fin clipping and PCR. We incrossed heterozygous fish for both lines and found that approximately 25% of their offspring showed the albino phenotype (data not shown), indicating that both attP insertions disrupted the *tyr* gene as expected. Together, these results indicate that we have established *tyr* mutant lines with the phiC31 landing site located in the pre-defined *tyr* loci.

**Table 2.**
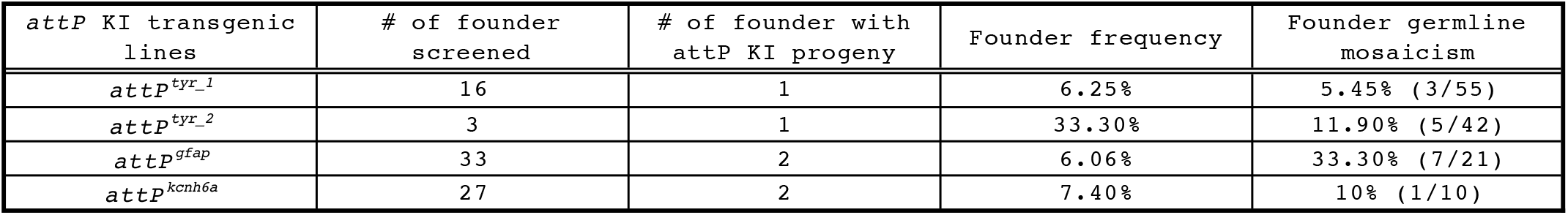
Founder frequencies and germline transmission mosaicisms of various attP knock-in (KI) alleles. For founder germline mosaicism, the results of one founder per target site are shown. The parentheses indicate the number of embryos containing the knock-in alleles over the number of embryos screened.

To evaluate the robustness of the attP knock-in method, we sought to insert attP into two more genes – the *glial fibrillary acidic protein (gfap)* gene and the *potassium voltage-gated channel, subfamily H, member 6a (kcnh6a)* gene. We designed and tested two to three gRNAs for each gene and identified one that yielded 57% mutation efficiency for *gfap* and another one that yielded 38.6% mutation efficiency for *kcnh6a* based on the PCR-fluorescent fragment length analysis (Supplemental Table S5). Next, we designed ssODNs to contain an attP encompassed by two short homology arms as described above, except that we avoided having a premature stop codon in the knock-in alleles of these genes (Supplemental Table S4). We performed microinjection of SpCas9 protein, gRNA, and ssODN, and we detected successful attP knock-ins in the microinjected embryos via next-generation sequencing (Table 1). Following similar genotyping strategies used for the *tyr* loci, we identified 2 *attP^gfap^* founders from 33 F_0_ fish screened and 2 *attP^kcnh6a^* founders from 27 F_0_ fish screened (Table 2). Thus, founder frequencies for *attP^gfap^* and *attP^kcnh6a^* were 6.1% and 7.4%, respectively. The knock-in sequences were verified (Figure 2G-H), and the germline mosaicism was 10% - 33% for the founders (Table 2). In sum, we have successfully created phiC31 transgenesis recipient lines in multiple zebrafish genes.

### Generation of allele-tracking reporter lines using phiC31 integrase

In theory, an attP site located in an endogenous gene will enable facile generation of a fluorescently tagged mutant allele via the phiC31 integrase technology, hence, eliminating the need for time-consuming genotyping procedures. To test this, we mated heterozygous *attP^tyr_1^* fish to the wild-type fish and microinjected their embryos with in vitro transcribed phiC31 mRNA together with the plasmid pDestattB_ubi:EGFP originally developed by Mosimann, et al (Mosimann *et al*., 2013b). This plasmid contains a 70-bp attB sequence as well as the *EGFP* gene driven by the *ubiquitin (ubi)* promoter, which can elicit ubiquitous green fluorescence from an early embryonic stage.(Mosimann et al., 2011) We used two sets of PCR primers to detect the 5’ and 3’ ends of phiC31-mediated integration in the injected embryos (Fig. 3A). The data showed that 13 out of 24 analyzed embryos exhibited correct integration at both 5’ and 3’ ends (Fig. 3B). Since it was expected that only half of the embryos would carry the attP knock-in allele, these results suggest that phiC31-mediated DNA recombination was very efficient at the *attP^tyr_1^* locus. We performed the same test using heterozygous *attP^tyr_2^* fish. However, we could not detect any integration events by PCR analysis, suggesting that the *attP^tyr_2^* locus may be defective or inaccessible for phiC31 integrase (data not shown). Consequently, we chose *attP^tyr_1^* for the following study. Sanger sequencing results confirmed the attL and attR sequences flanking the integration of pDestattB_ubi:EGFP at the *attP^tyr_1^* locus, indicating precise recombination between attB and attP (Fig. 3C). Taken together, these results demonstrate that phiC31 integrase can mediate precise and efficient DNA integration in zebrafish.

**Figure 3.**
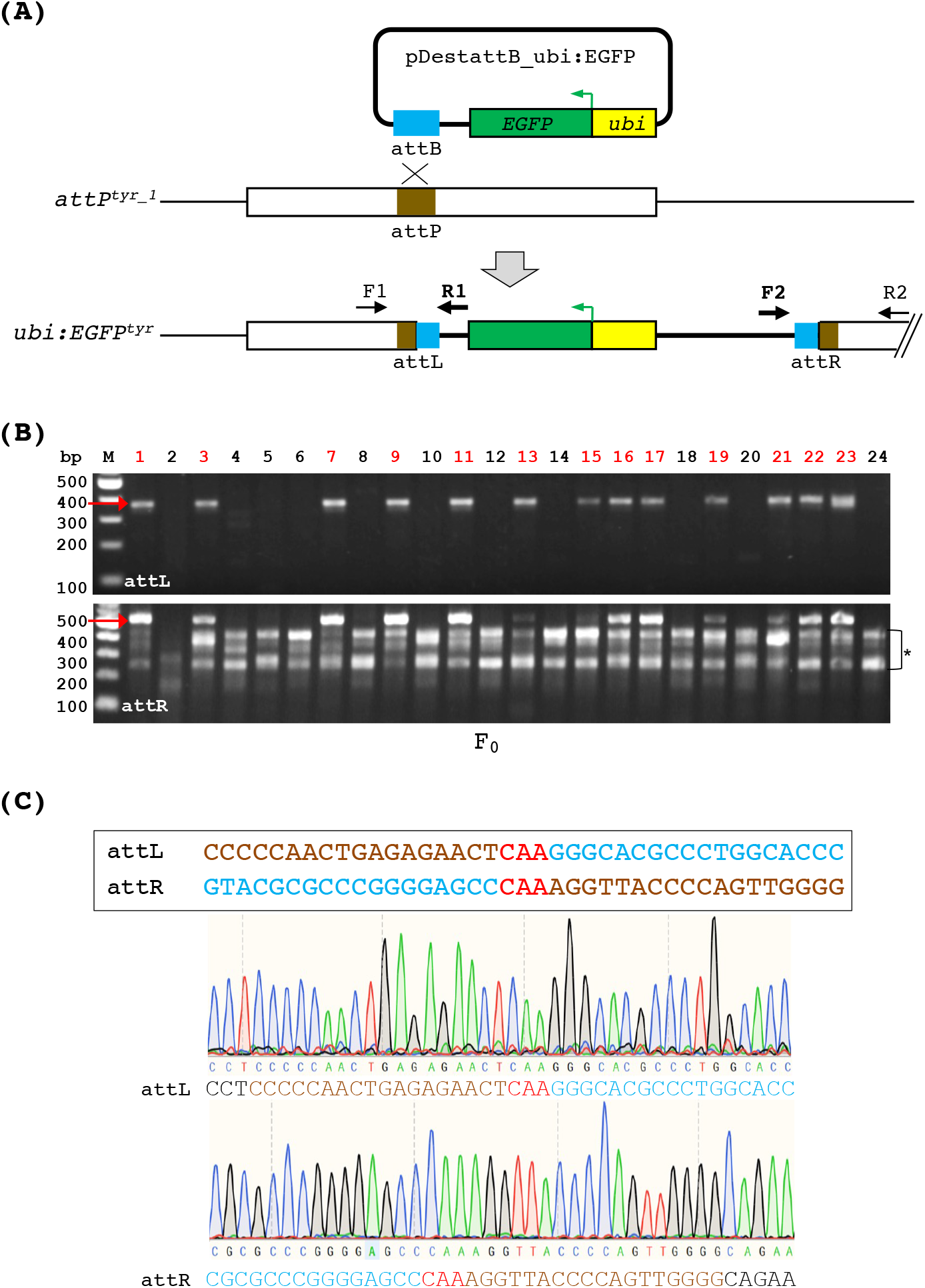
PhiC31 integrase mediates DNA integration via recombination between attB and the genomic attP site. (A) A diagram depicting the integration of *ubi:EGFP* at the *attP^tyr_1^* locus. PhiC31 integrase mediates recombination between the plasmid DNA containing the attB sequence and the genomic attP site, resulting in single-copy integration of the entire plasmid. The integrated DNA (thick black line) is flanked by two attP/attB composite sequences named attL and attR, which can be detected via PCR using two sets of primers (F1/R1 and F2/R2) as indicated. F1 and R2 are *tyr*-specific, whereas R1 and F2 are plasmid-specific. (B) PCR analysis of attL and attR in individual embryos microinjected with the phiC31 mRNA and pDestattB_ubi:EGFP. Red arrows indicate the expected product sizes of attL (369 bps, top panel) and attR (534 bps, bottom panel). The bracket with an asterisk on the right side of the bottom gel indicates non-specific amplifications from the F2/R2 primer set. The embryos with the correct size products from both attL and attR PCR are indicated by red numbers on the top. (C) Sanger sequencing of the attL (top chromatogram) and attR (bottom chromatogram) PCR products demonstrates precise DNA integration. Reference sequences for attL and attR are shown in the box above the chromatograms. Sequences originating from attP and attB are indicated in brown and turquoise, respectively. The ‘CAA’ crossover site is shown in red.

To evaluate germline transmission of the recombinant allele, we raised the injected embryos from heterozygous *attP^tyr_1^* outcrosses to adulthood. We screened three fish and identified one founder that produced green-fluorescent offspring (Figure 4A). The ratio between green and non-green F_1_ embryos was approximately 1:1. Using PCR, we could detect the correct integration of *ubi:EGFP* at the *attP^tyr_1^* locus in 50% of the green fluorescent embryos, suggesting that this founder carried not only phiC31-mediated but also random integration (Figure 4B). We did not investigate the location of the off-target integration in this founder. Presently, there are no known functional pseudo-attP sites in the zebrafish genome.

**Figure 4.**
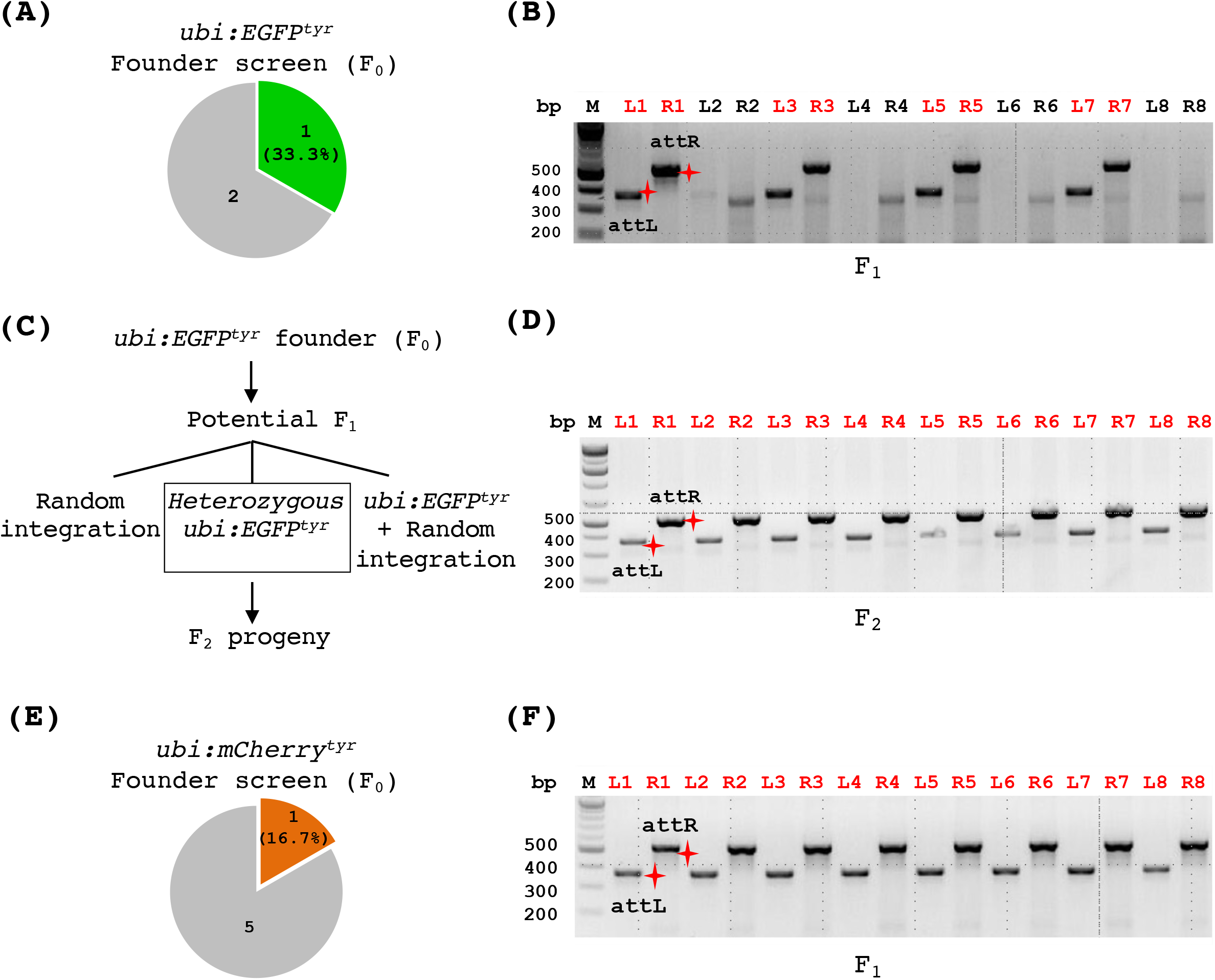
Generation of allele-tracking reporter lines for the *tyr* mutation. (A-B) Founder screen results of the transgenic *ubi:EGFP^tyr^* zebrafish. (A) A pie chart indicating the number of F_0_ fish screened that produced some fluorescent embryos (the green piece) or all non-fluorescent embryos (the gray piece). (B) PCR analysis of individual fluorescent F_1_ embryos demonstrating that some of them (labeled by red numbers on top) carried a correct *ubi:EGFP* integration. ‘L’ and ‘R’ in front of the numbers on top indicate attL and attR PCR, respectively. The red shining star symbols mark the correct size products of attL (369 bps) and attR (534 bps). (C) Schematic depicting potential genotypes of the descendants from an *ubi:EGFP^tyr^* founder fish. (D) PCR analysis of individual fluorescent embryos (F_2_) from a heterozygous *ubi:EGFP^tyr^* fish (F_1_) outcrossed to a wild-type fish indicating that all of them carried the correct integration. (E-F) Founder screen results of the transgenic *ubi:mCherry^tyr^* zebrafish. (E) A pie chart indicating the number of F_0_ fish screened that produced some fluorescent embryos (the red piece) or all non-fluorescent embryos (the gray piece). (F) PCR analysis of individual fluorescent F_1_ embryos showing that all of them carried a correct *ubi:mCherry* integration.

Moreover, it has been shown that random plasmid integration occurs less frequently compared to phiC31-mediated integration into transgenic attP sites in zebrafish.(Mosimann *et al*., 2013b) We raised green-fluorescent F_1_ fish to adulthood and performed fin-clipping and PCR to identify the F_1_ fish that harbored the correct *ubi:EGFP* integration. Since these fish could still carry more than one *EGFP* integration (Figure 4C), we crossed 8 fish carrying the correct integration to the wild-type fish and found that 5 out of 8 fish produced approximately 1:1 of green versus non-green embryos. To confirm that these F1 fish did not carry random integration, we conducted PCR analysis of the F2 progeny and found that all fluorescent embryos we analyzed had a correct integration (Figure 4D). Thus, these results indicate that we have successfully generated heterozygous fish that carry a single-copy, *ubi:EGFP* reporter in the *tyr* gene (denoted as the *ubi:EGFP^tyr^* allele).

We carried out a similar workflow to generate a red fluorescent report line to track the *tyr* mutant allele. We generated the pDestattB_ubi:mCherry construct and injected it along with the phiC31 mRNA into the embryos of heterozygous *attP^tyr_1^* fish crossed to the wild-type fish. When the injected embryos reached adulthood, we screened six F_0_ fish that exhibited high mCherry mosaicism and identified one founder (Figure 4E). This founder fish produced approximately 50% red fluorescent progeny when outcrossed to a wild-type fish. Moreover, all fluorescent embryos analyzed by PCR showed a correct integration (Figure 4F). We raised red fluorescent F_1_ fish to adulthood, outcrossed 3 fish to the wild-type fish, and found that all 3 fish carried a single-copy integration of *ubi:mCherry* at the *tyr* gene. Overall, these results demonstrate that attP fish lines are useful transgenesis recipients. One attP fish line can be used to derive multiple integration lines with ease. Meanwhile, phiC31 integrase can mediate efficient and transmissible DNA integration at genomic attP landing sites.

### Instantaneous visual genotyping using the *tyr* allele-tracking reporter lines

Having created the allele-tracking reporter lines with two different colors for the *tyr* gene, we sought to demonstrate the feasibility of a technique for instantaneous visual genotyping. In this method, by mating two heterozygous mutant fish that each has its own marker linked to the same mutation, one can determine whether a progeny is a wild type, heterozygote, or homozygote simply by examining its color. Hence, we mated a heterozygous *ubi:EGFP^tyr^* fish to a heterozygous *ubi:mCherry^tyr^* fish and found that all double fluorescent fish exhibited the albino phenotype and vice versa (Fig. 5). Thus, these results demonstrate the advantages of constructing mutant lines along with allele-tracking reporters via the combination of CRISPR-Cas9 and phiC31 technologies.

**Figure 5.**
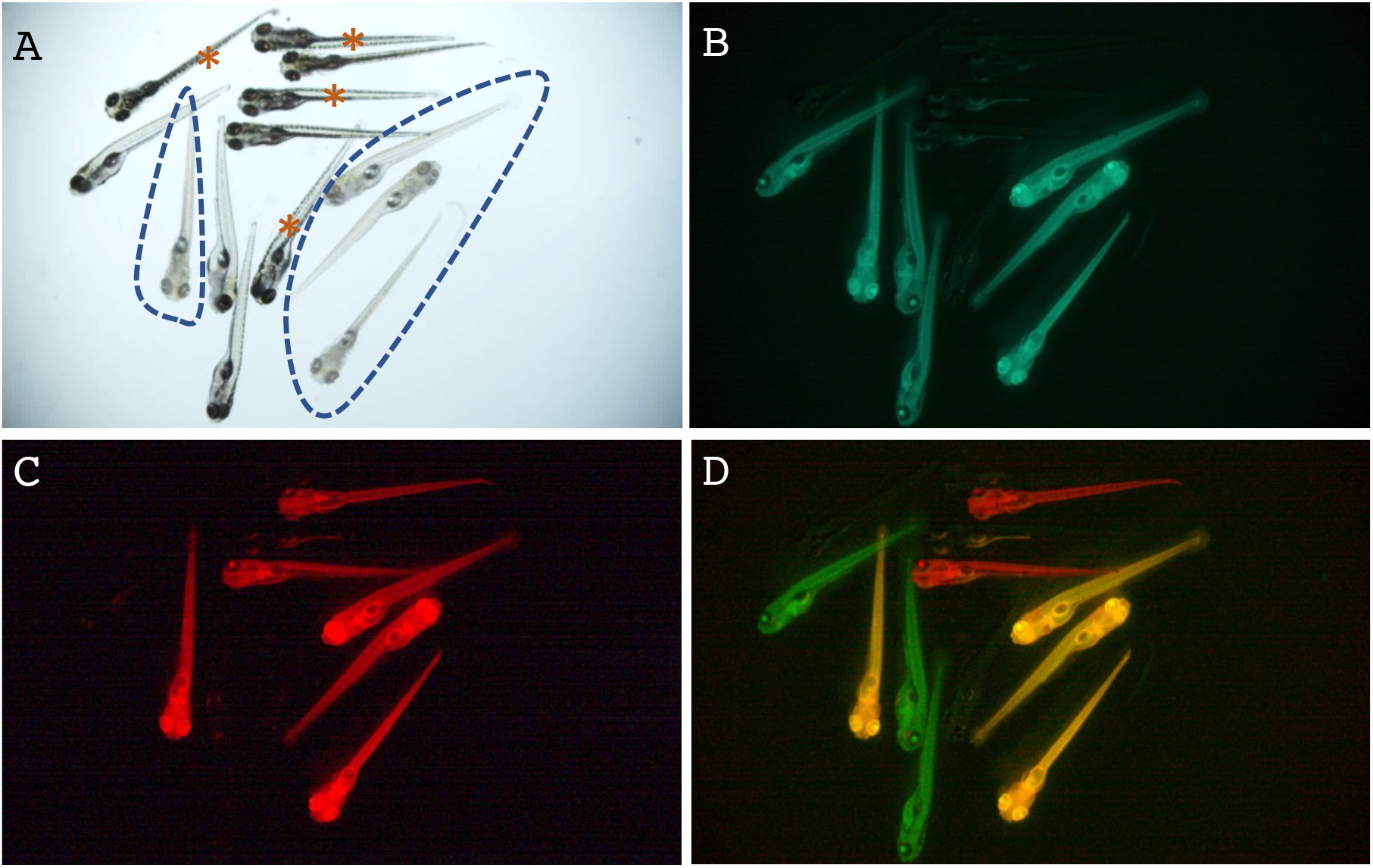
Instantaneous visual genotyping using the *tyr* allele-tracking reporter lines. (A-D) Visual genotyping results of the *tyr* mutation using *ubi:EGFP^tyr^* and *ubi:mCherry^tyr^*, two allele-tracking reporter lines. Heterozygous fish of these two lines were intercrossed, and their offspring were inspected under a fluorescent microscope. The same group of embryos was imaged in the bright field (A), green-fluorescent channel (B), and red-fluorescent channel (C). Images of (B) and (C) were merged in (D). In panel A, embryos exhibiting the homozygous *tyr* mutant phenotype as evidenced by the loss of pigmentation were encircled by dash lines. In panel D, embryos that were both green and red were shown in yellow. Embryos that had no fluorescence in (B-D) were indicated by brown asterisks in (A). Hence, the wild-type embryos were unmarked, the heterozygous embryos were either green or red, and the homozygous embryos were both green and red.

### Generation of a target gene-specific reporter line via TICIT

Another potential application of single-copy transgenesis at a user-specified genomic location is to generate gene-tagging or target gene-specific reporter lines. As a proof of concept, we sought to insert a reporter gene into the *attP^gfap^* locus. During embryonic development, *gfap* is abundantly expressed in the glial cells of the eye and the central nervous system (CNS). To do this, we first generated a plasmid pDestattB_P2A-EGFP by inserting the coding sequences of the self-cleaving peptide P2A and a promoter-less *EGFP* gene next to the attB site in pDestattB (Addgene #68313). We expected that phiC31-mediated recombination between the plasmid DNA and the genomic *attP^gfap^* locus should result in an in-frame integration of *EGFP* after the start codon of *gfap* (Figure 6A). Thus, we microinjected pDestattB_P2A-EGFP and the phiC31 mRNA into the embryos of heterozygous *attP^gfap^* fish outcrossed to the wild-type fish. In the injected embryos, we could readily see mosaic *EGFP* expression specifically in the CNS (Figure 6B). When the injected embryos reached adulthood, we screened one F_0_ fish with *EGFP* expression in the CNS and found that it produced 18% fluorescent progeny (32 out of 175 embryos). All F_1_ fluorescent embryos expressed a consistent level and pattern of *EGFP* expression in the CNS (Figure 6B), and all fluorescent embryos analyzed by PCR showed a correct integration (Figure 6C). Further, confocal imaging analysis showed EGFP expression in the eye and the brain one-day post fertilization (dpf) (Figure 6D). While the fluorescence in the head region subsided after 1 dpf, its intensity in the posterior CNS region persisted through later stages, correlating with the reported *gfap* expression pattern (https://zfin.org). These results demonstrate that TICIT can be applied to the generation of transgenic animals in which transgene expression is controlled by an endogenous promoters of interest.

**Figure 6.**
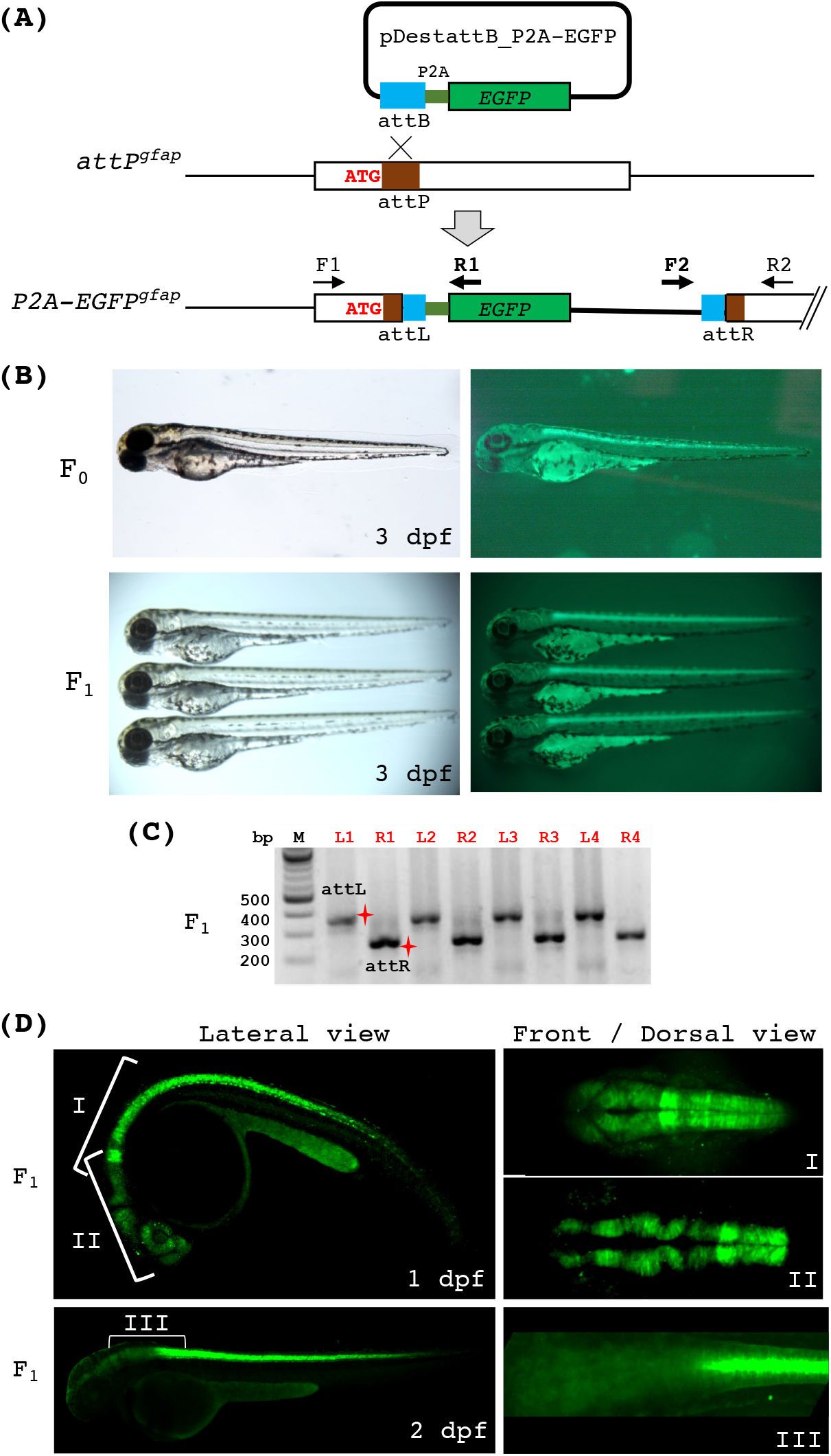
Generation of the endogenous *gfap* reporter line via phiC31 integrase. (A) A diagram depicting the integration of a promoter-less EGFP reporter at the *attP^gfap^* locus via phiC31-mediated recombination. Correct integration of pDestattB_P2A-EGFP will result in fusion and co-translation of the N-terminus of gfap and P2A-EGFP. We named this allele *P2A-EGFP^gfap^*. Two sets of PCR primers (F1/R1 and F2/R2 as indicated) were used to detect the integration. F1 and R2 are *gfap*-specific, whereas R1 and F2 are plasmid-specific. (B) Bright-field and fluorescence images of endogenous *gfap* reporter fish embryos at 3 dpf. The top panels show an *attP^gfap^* embryo after microinjection with the phiC31 mRNA and pDestattB_P2A-EGFP. The bottom panels show three heterozygous *P2A-EGFP^gfap^* embryos. (C) PCR analysis of individual fluorescent embryos from a founder fish indicating that all of them harbored the correct integration. ‘L’ and ‘R’ in front of the numbers on top indicate attL and attR PCR, respectively. The red shining star symbols mark the correct size products of attL (332 bps) and attR (282 bps). (D) Stacked confocal images of the *P2A-EGFP^gfap^* embryos showing various regions and angles of view at 1 and 2 dpf.

### The *attP^tyr^* landing site supports transgene expression in a wide range of tissue and cell types

In the *ubi:EGFP^tyr^* fish, we noticed broad and strong green fluorescence from an early embryonic stage, suggesting that the *attP^tyr^* landing site may act as a safe harbor locus for transgene expression. To investigate this further, we examined the fluorescence of heterozygous *ubi:EGFP^tyr^* fish from embryonic to adult stages. Using a fluorescent stereoscope, we could see faint EGFP expression starting from 6 hours post fertilization (hpf) (Figure 7A). The intensity of green fluorescence soon became clear, appeared ubiquitous, and continued throughout the embryonic stages (Fig. 7A). All heterozygous *ubi:EGFP^tyr^* adult fish showed consistently strong and ubiquitous green fluorescence from outside. We dissected the adult fish and found that *EGFP* was expressed in all organs and tissue types, such as muscle, gill, eye, brain, skin, spleen, heart, liver, kidney, intestine, pancreas, testis, and ovary (Supplemental Figure S1A). However, the fluorescence was noticeably absent in mature eggs, which was different from another *ubi* reporter line *(ubi:loxP-EGFP-loxP-mCherry*, also known as *ubi:Switch*) that showed maternally deposited *EGFP* expression (Supplemental Figure S1B) (Mosimann *et al*., 2011). Next, to get a closer look at some of the tissue and cell types, we performed immunohistochemistry (IHC) using an anti-GFP antibody. The results showed that EGFP could be detected in all tissue sections, even though it was not at the same level among different cell types (Figure 7B). These data are expected as it was previously shown that the *ubi* promoter renders a ‘ubiquitous’, but not necessarily ‘homogenous’ expression in all cell types (Mosimann *et al*., 2011). Further, we isolated hematopoietic cells from the whole kidney marrow, which is the hemogenic tissue in zebrafish, and analyzed the fluorescence of various blood lineages using flow cytometry (Figure 7C). We found that *EGFP* was expressed in >90% of hematopoietic progenitors, lymphocytes, and myelomonocytes and in ~40% of erythrocytes. Taken together, these findings demonstrate that the *attP^tyr^* landing site can support transgene expression in almost all tissue and cell types. Accordingly, site-specific transgenesis using the *attP^tyr^* locus may be advantageous compared to Tol2-based transgenesis because it can circumvent positional effects and yield predictable and reproducible transgene expression as controlled by its own promoter.

**Figure 7.**
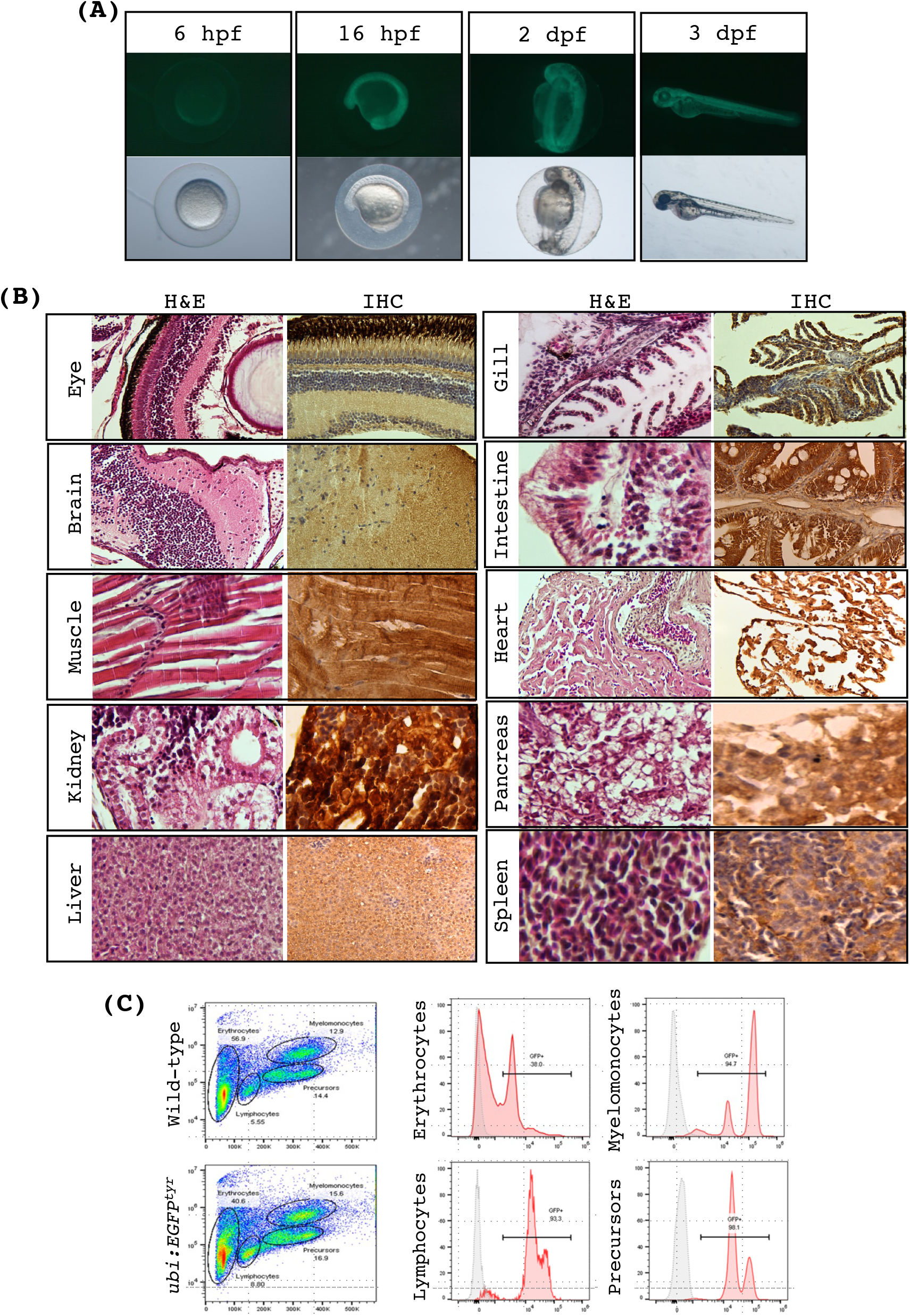
*ubi:EGFP^tyr^* transgenic fish exhibit robust and ubiquitous EGFP expression. (A) Bright-field and fluorescence images of *ubi:EGFP^tyr^* embryos at various developmental stages as indicated. hpf, hours post fertilization; dpf, days post fertilization. (B) Histology sections of *ubi:EGFP^tyr^* adult fish stained with H&E or anti-EGFP antibody (IHC) demonstrating widespread but varying levels of EGFP expression among different tissue and cell types. (C) Representative flow cytometry analysis of the wild-type and *ubi:EGFP^tyr^* adult whole kidney marrow (WKM). Major blood lineages in WKM were resolved and identified as previously shown (left panels).^33^ EGFP expression was analyzed in each blood cell gate (middle and right panels). Gray, wild-type; pink, *ubi:EGFP^tyr^*.

## Discussion

The development of research tools can serve as a potent catalyst for biomedical research. Genome engineering tools have been profound examples. In recent years, CRISPR technologies have been widely embraced by the scientific community and have spawned tens of thousands of publications and applications (Anzalone et al., 2020). However, even with current CRISPR methods, the efficiencies for targeted integration of large DNA fragments in mammalian cells and various model organisms remain low, largely due to their dependencies on cellular DNA repair mechanisms (Prill and Dawson, 2020; Yang et al., 2020). In contrast, phiC31 integrase mediates DNA integration independently and with high efficiencies (Mosimann *et al*., 2013b; Roberts *et al*., 2014). However, their recognition sites are not programmable. In this study, we demonstrate that targeted integration of long DNA fragments can be achieved via a 2-step protocol. First, a short phiC31 attachment sequence is inserted into a selected genomic target site by CRISPR-Cas9. This will enable subsequent DNA integration into the same target site via phiC31. A workflow of this method is provided in Figure S2. As summarized in Figure 1, we show that this new technique, named TICIT, can open doors to diverse downstream applications for zebrafish research.

In the first step of this protocol, we successfully inserted the attP landing pad into various targeted loci in a homology-dependent manner via SpCas9 and ssODNs at a founder frequency ranging between 6-33% (Table 2). This method is simple and effective, and it does not require any DNA cloning. Moreover, using this method, researchers can place the attP site precisely in a way that either creates or avoids an in-frame stop codon for different applications, as we have shown in this report (Figure 2).

In the second step of this protocol, we only needed to screen a small number of fish to identify a founder. The founder frequency ranged between 16-100% at two sites — attP^tyr_1^ and attP^gfap^. However, we could not detect any phiC31-mediated recombination at the attP^tyr_2^ site. Defective attP landing sites have also been reported previously (Mosimann *et al*., 2013b). Presently, it is unknown why some attP landing sites are refractory to phiC31 integrase. Nonetheless, we showed that zebrafish lines carrying a functional landing site can be used repetitively with high efficiency to generate multiple integration lines. With this method, we developed technology for marking zebrafish lines with fluorescent reporters indicating whether the line is wild-type, heterozygous, or homozygous for an allele of interest. This technique has the potential to be incredibly valuable for routine zebrafish work, allowing genotyping by visual inspection, thereby saving enormous amounts of time and money. It could be equally valuable for researchers working with mice, rats, or other organisms. The technique may also enable new experiments by allowing early identification of mutant animals from a mixed population. For example, it can be used to sort a homogeneous population of homozygous mutants for use in high-throughput screening for potential treatments of genetic disorders. Here, we used the *ubi* promoter to enable early detection and easy sorting. However, researchers can choose any promoters that better suit their purposes. Furthermore, the results showed that phiC31 mediated unidirectional integration with high fidelity, since we could identify correct attL and attR sequences flanking the inserted DNA. This enables efficient in-frame insertion of a fluorescent marker into a targeted gene, which may prove useful when studying previously uncharacterized genes. In addition to reporter genes, one can also express other transgenes from an endogenous promoter. For example, by expressing the cDNAs of various genetic variants, this technique could be used for gene replacement in comparative studies of genetic variations.

One of the greatest benefits of single-copy, site-specific transgenesis as compared to transgenesis via random integration is the ability to obtain faithful transgene expression controlled by its own promoter without positional and copy number artifacts. This could be achieved by using phiC31 integrase and a zebrafish line that carries a ‘safe harbor’ phiC31 docking locus. Hence, the development of this technique may widen the use of the zebrafish as a platform to rapidly assess the functions of genetic variants in the coding and non-coding regions identified in genome-wide association studies (GWAS) (Edwards et al., 2013; Gusev et al., 2018; Tucker et al., 2017). Indeed, the phiC31 transgenesis method via several Tol2-generated attP lines has been shown to be superior to the Tol2 transgenesis method when evaluating human cis-regulatory elements in zebrafish (Bhatia et al., 2021; Roberts *et al*., 2014). The landing sites in the integration recipient lines used in previous studies have been mapped to both intergenic and intragenic regions (Bhatia *et al*., 2021; Mosimann *et al*., 2013b; Roberts *et al*., 2014). Here, we demonstrate that the *attP^tyr_1^* landing site exhibit high recombination efficiency (Figure 4). Fish that harbor *ubi:EGFP* and *ubi:mCherry* in the *attP^tyr_1^*locus showed strong, ubiquitous, and consistent levels of expression over several generations (Figure 7 and Supplemental Figure S1). Though transgene integration at the *attP^tyr_1^* site will be accompanied by a single-copy deletion of the *tyr* gene, we have not observed any noticeable phenotypes in heterozygous *tyr* zebrafish, suggesting that the *attP^tyr_1^* fish can potentially be used as neutral transgenesis recipients.

Taken together, we show that CRISPR-Cas9 and phiC31 technologies can be efficiently combined to construct novel genome engineering tools and zebrafish models. Presumably, it may be possible to perform attP insertion and DNA integration via phiC31 in one single step, which has been successfully demonstrated in human cells using the prime editor system instead of SpCas9 (Anzalone et al., 2022). Thus, this will be a useful future direction for zebrafish. Moreover, it will be helpful to identify other microbial DNA integrases that exhibit high efficiencies in zebrafish, which will enable more sophisticated experimental designs involving multiple DNA integrations or cassette exchange (Durrant, 2021; Jusiak et al., 2019; Low et al., 2022). It can be expected that a broader adoption and more creative uses of the CRISPR and integrase technologies in zebrafish and other model organisms will play an important role in accelerating more transformative biomedical research in the near future.

## Acknowledgments

This work was supported by the Hassenfeld Scholar Award (to J.-R. J. Yeh) and NIH no. R01 GM134069 (to R. T. Peterson and J.-R. J. Yeh). J. Ma received support from the China Scholarship Council (no. 201808210354).

## Author contributions

J. Ma, R.T. Peterson, and J.-R. J. Yeh conceived and designed the research; J. Ma, W. Zhang, Z. Sun, and S. Parvez performed the research and acquired the data, J. Ma, W. Zhang, and Z. Sun analyzed and interpreted the data. J. Ma and J.-R. J. Yeh drafted the manuscript. All authors were involved in reviewing and revising the manuscript.

---

**Figure S1.**
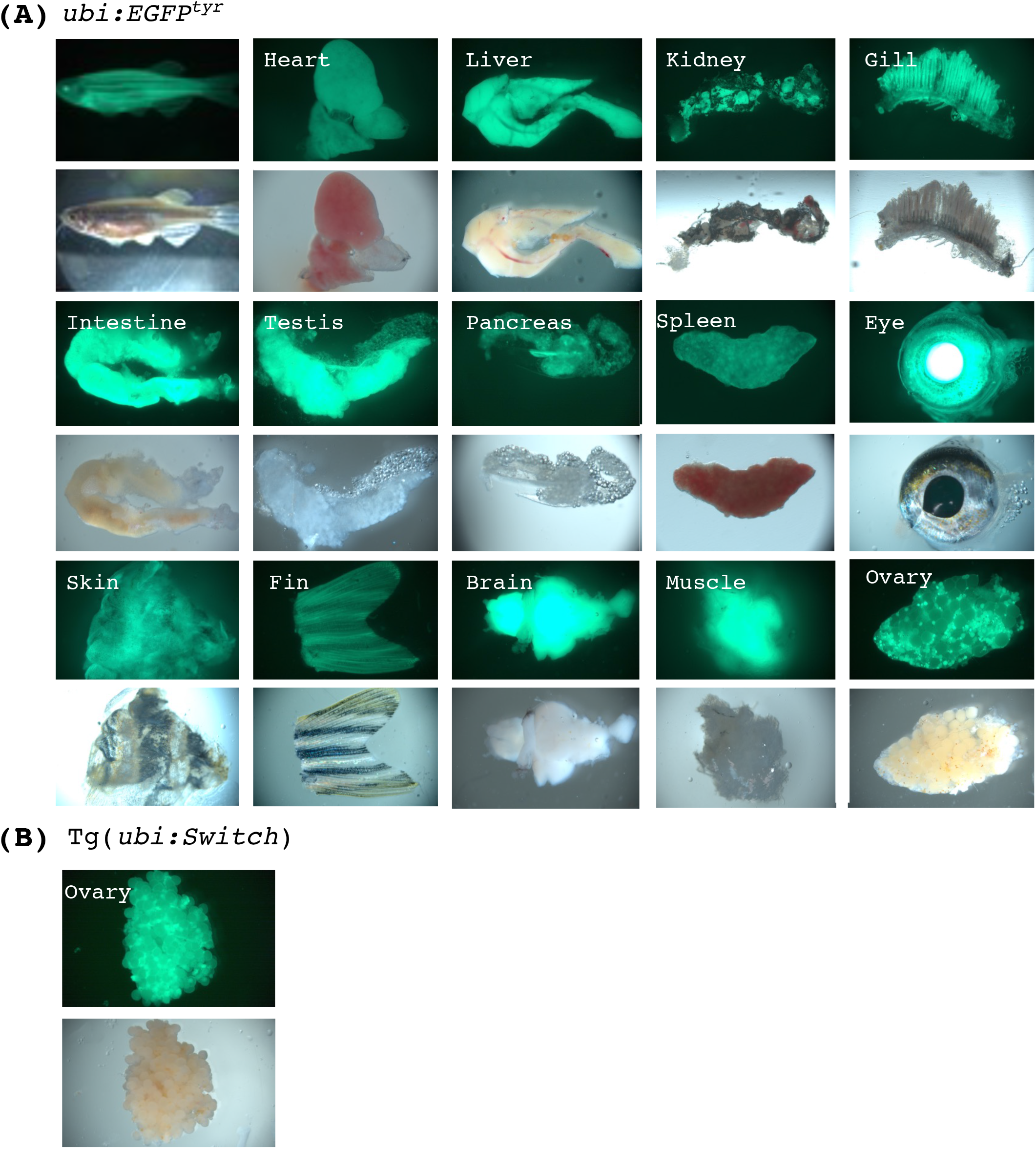
EGFP fluorescence can be seen in multiple organs and tissues of *ubi:EGFP^tyr^* adult fish. (A) Bright-field and fluorescence images of various organs and tissues isolated from *ubi:EGFP^tyr^* adult female fish. Broad EGFP expression patterns are detected. Note the fluorescence in the ovary indicates that EGFP expression can be detected in immature oocytes but not in mature oocytes. (B) Bright-field and fluorescence images of the ovary from *Tg(ubi:Switch)* adult female fish. Note that EGFP expression can be detected in both immature and mature oocytes.

**Figure S2.**
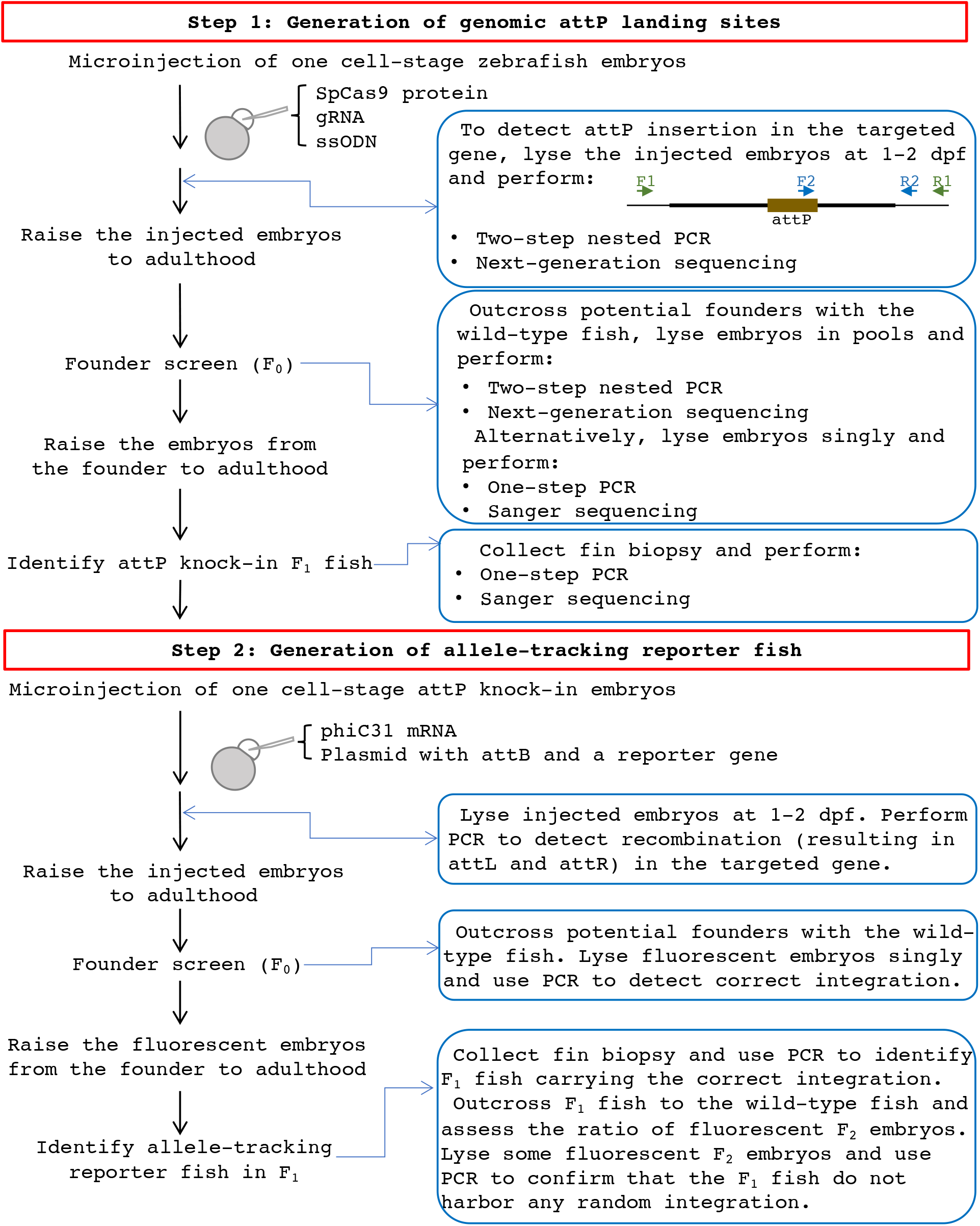
The workflow for the generation of allele-tracking reporter zebrafish using TICIT. The flowchart outlines the procedures for generating allele-tracking reporter zebrafish.

**Table S1.**
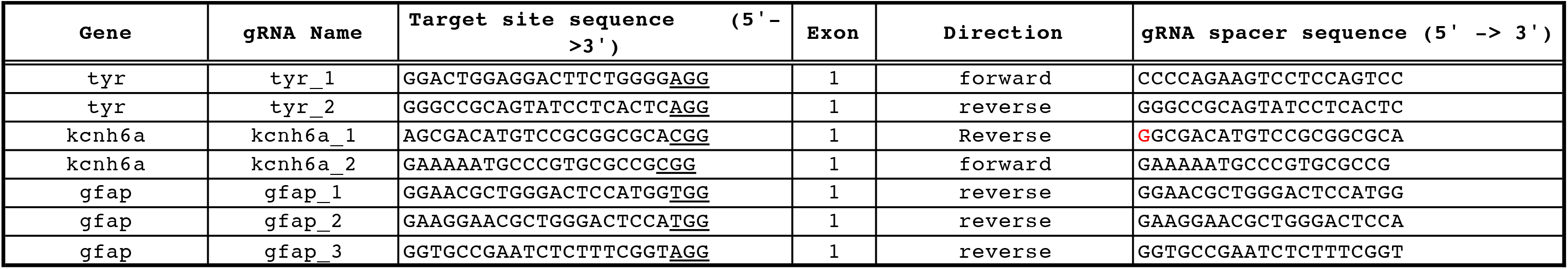
SpCas9 target sites tested or used in this study. In the ‘Target site sequence’, PAM sequence is underlined. In the ‘gRNA spacer sequence’, an extra ‘G’ (shown in red) is added to faciliate efficient in vitro transcription. For attP knock-in, kcnh6a 2 and gfap 1 gRNAs were used.

**Table S2.**
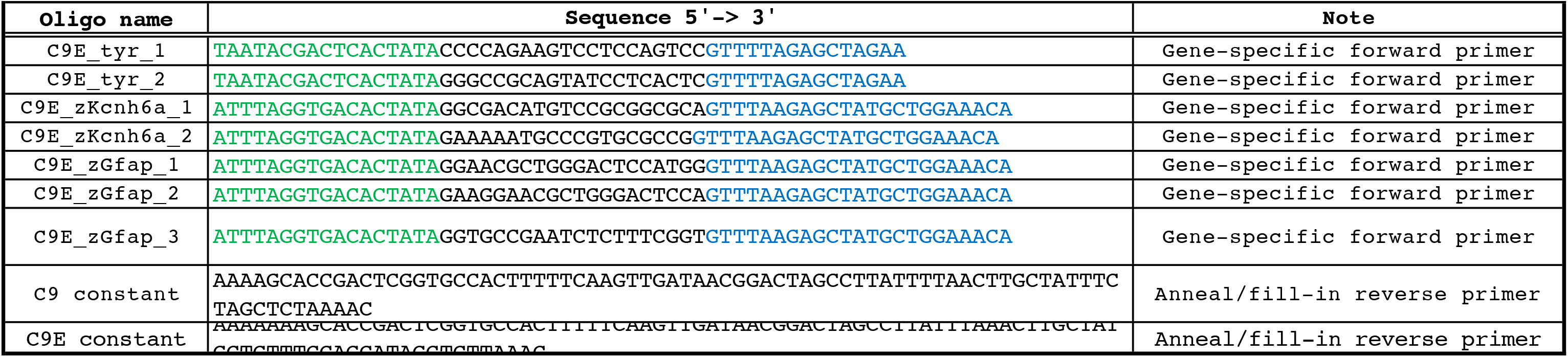
Oligonucleotides used for gRNA construction. In ‘Sequnce’, T7 or SP6 promoter sequences are shown in green, and the sequences complementary to the C9 or C9E constant oligo are shown in blue.

**Table S3.**
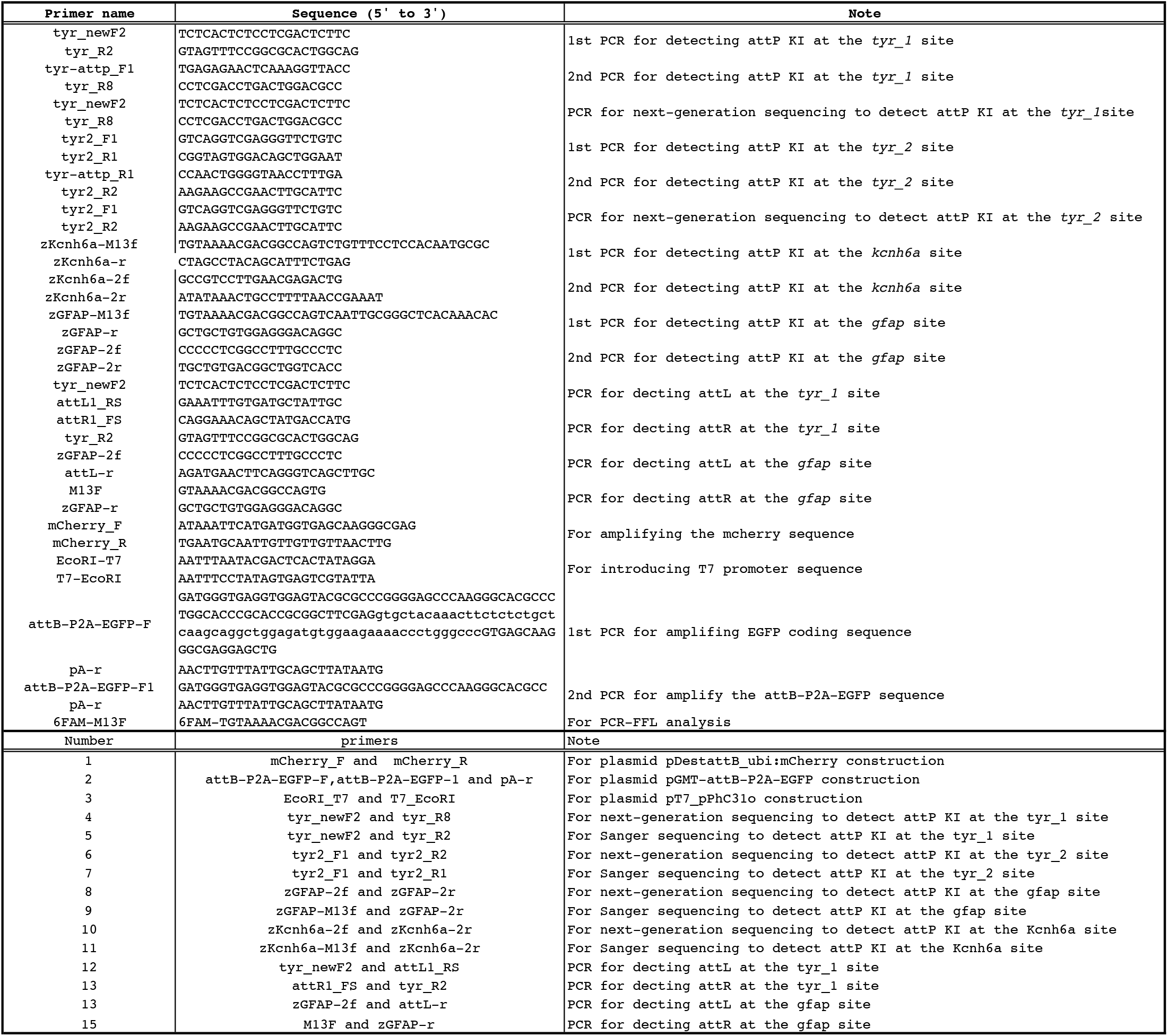
PCR and cloning primers used in this study.

**Table S4.**
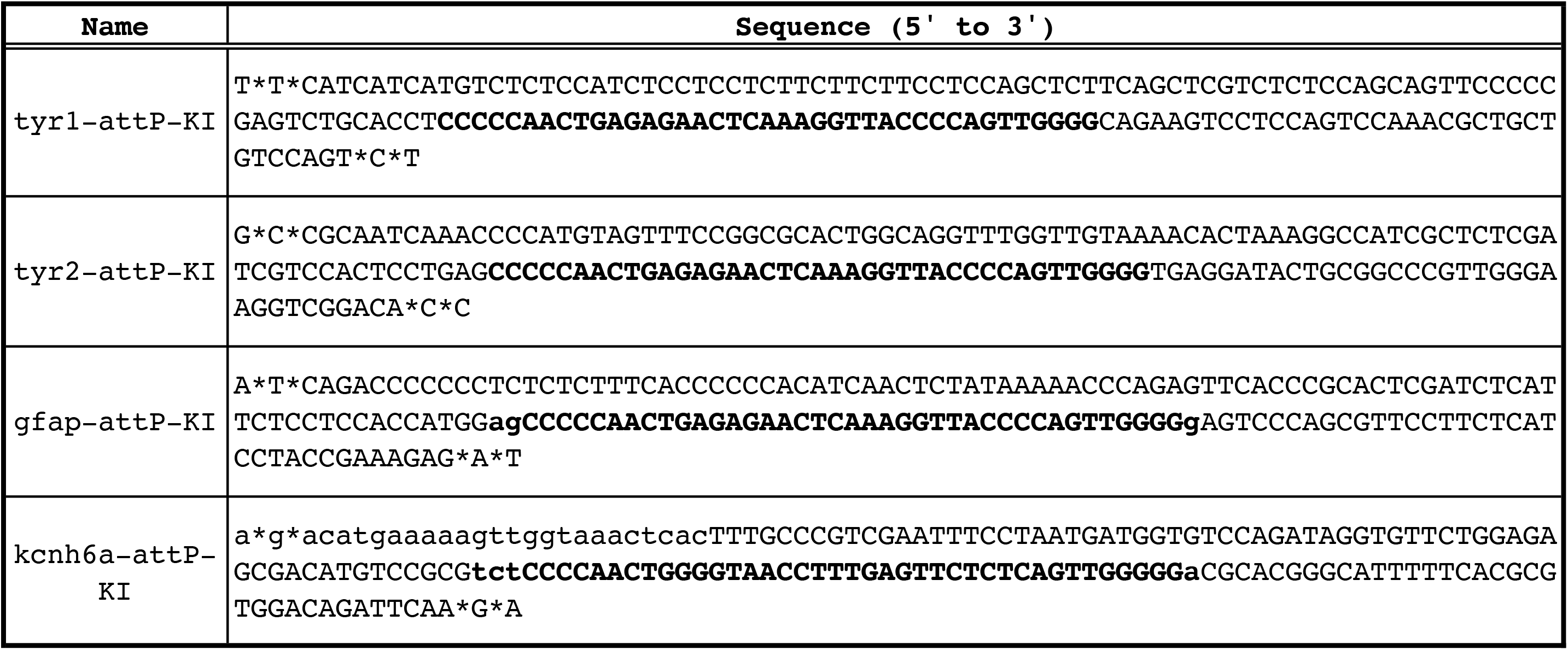
Sequences of the single-stranded oligonucleotides (ssODNs) used for attP knockin experiments. The ssODNs were chemically synthesized and two phosphorothioate linkages (denoted by asterisks) were added to both termini to enhance stability. The knock-in sequences are shown in bold. The attP sequences are shown in bold uppercase letters. The bold lowercase letters were added to avoid stop codons in the reading frames.

**Table S5.** Fluorescent PCR fragment length analysis for gRNA efficiency. For each gRNA, two pools of embryos (10 embryos per pool) were analyzed. Samples K5, K6, G7, and G8 are uninjected control embryos. PCR products were run on ABI 3730xl DNA Analyzer at MGH DNA core. Fragment sizes, peak heights, and peak areas were determined using GeneMapper v4.0. The fragments correspond to the wild-type alleles are highlighted in yellow. gRNA efficiencies were determined by comparing the abundance of the wild-type alleles over the sum of all peaks in the same samples based on peak intensity. For attP knock-in, kcnh6a_2 and gfap_1 gRNAs were used.

| Sample Name | gRNA | Size 1 | Size 2 | Size 3 | Size 4 | Size 5 | Size 6 | Size 7 | Size 8 | Size 9 | Height 1 | Height 2 | Height 3 | Height 4 | Height 5 | Height 6 | Height 7 | Height 8 | Height 9 | Peak Area 1 | Peak Area 2 | Peak Area 3 | Peak Area 4 | Peak Area 5 | Peak Area 6 | Peak Area 7 | Peak Area 8 | Peak Area 9 |
| --- | --- | --- | --- | --- | --- | --- | --- | --- | --- | --- | --- | --- | --- | --- | --- | --- | --- | --- | --- | --- | --- | --- | --- | --- | --- | --- | --- | --- |
| K1 | kcnh6a_1 | 286.59 |  |  |  |  |  |  |  |  | 9269 |  |  |  |  |  |  |  |  | 105184 |  |  |  |  |  |  |  |  |
| K2 | kcnh6a_1 | 286.59 |  |  |  |  |  |  |  |  | 10777 |  |  |  |  |  |  |  |  | 120346 |  |  |  |  |  |  |  |  |
| K3 | kcnh6a_2 | 280.17 | 282.13 | 286.68 |  |  |  |  |  |  | 1242 | 862 | 4446 |  |  |  |  |  |  | 12559 | 8309 | 47710 |  |  |  |  |  |  |
| K4 | kcnh6a_2 | 280.22 | 282.12 | 286.65 |  |  |  |  |  |  | 2021 | 1456 | 3728 |  |  |  |  |  |  | 20934 | 13569 | 39176 |  |  |  |  |  |  |
| K5 | - | 286.71 |  |  |  |  |  |  |  |  | 9192 |  |  |  |  |  |  |  |  | 100197 |  |  |  |  |  |  |  |  |
| K6 | - | 286.69 |  |  |  |  |  |  |  |  | 10088 |  |  |  |  |  |  |  |  | 111595 |  |  |  |  |  |  |  |  |
| G1 | gfap_1 | 246.18 | 254.89 | 264.28 |  |  |  |  |  |  | 127 | 302 | 1152 |  |  |  |  |  |  | 1437 | 2886 | 11533 |  |  |  |  |  |  |
| G2 | gfap_1 | 243.03 | 255.05 | 257.96 | 260.66 | 264.44 | 280.25 |  |  |  | 80 | 659 | 186 | 757 | 323 | 167 |  |  |  | 700 | 6968 | 1669 | 7948 | 2903 | 1798 |  |  |  |
| G3 | gfap_2 | 235.1 | 238.64 | 250.11 | 252.89 | 254.82 | 257.6 | 262.3 | 265.07 | 272.9 | 62 | 74 | 240 | 305 | 256 | 115 | 582 | 91 | 68 | 1010 | 592 | 2493 | 3288 | 2405 | 995 | 6105 | 951 | 1021 |
| G4 | gfap_2 | 231.08 | 242.96 | 250 | 254.67 | 261.2 | 264.95 | 269.75 | 278.18 |  | 212 | 646 | 178 | 981 | 591 | 308 | 221 | 362 |  | 2207 | 6682 | 1775 | 10592 | 6482 | 3157 | 2158 | 4126 |  |
| G5 | gfap_3 | 263.99 |  |  |  |  |  |  |  |  | 4402 |  |  |  |  |  |  |  |  | 47518 |  |  |  |  |  |  |  |  |
| G6 | gfap_3 | 264 |  |  |  |  |  |  |  |  | 4310 |  |  |  |  |  |  |  |  | 46587 |  |  |  |  |  |  |  |  |
| G7 | - | 264.08 |  |  |  |  |  |  |  |  | 5391 |  |  |  |  |  |  |  |  | 61207 |  |  |  |  |  |  |  |  |
| G8 | - | 264.02 |  |  |  |  |  |  |  |  | 5805 |  |  |  |  |  |  |  |  | 66142 |  |  |  |  |  |  |  |  |

## References

Abe, G., Suster, M.L., and Kawakami, K. (2011). Tol2-mediated transgenesis, gene trapping, enhancer trapping, and the Gal4-UAS system. Methods Cell Biol 104, 23–49. 10.1016/B978-0-12-374814-0.00002-1.

Anzalone, A.V., Gao, X.D., Podracky, C.J., Nelson, A.T., Koblan, L.W., Raguram, A., Levy, J.M., Mercer, J.A.M., and Liu, D.R. (2022). Programmable deletion, replacement, integration and inversion of large DNA sequences with twin prime editing. Nat Biotechnol 40, 731–740. 10.1038/s41587-021-01133-w.

Anzalone, A.V., Koblan, L.W., and Liu, D.R. (2020). Genome editing with CRISPR-Cas nucleases, base editors, transposases and prime editors. Nat Biotechnol 38, 824–844. 10.1038/s41587-020-0561-9.

Auer, T.O., Duroure, K., De Cian, A., Concordet, J.P., and Del Bene, F. (2013). Highly efficient CRISPR/Cas9-mediated knock-in in zebrafish by homology-independent DNA repair. Genome research. 10.1101/gr.161638.113.

Belteki, G., Gertsenstein, M., Ow, D.W., and Nagy, A. (2003). Site-specific cassette exchange and germline transmission with mouse ES cells expressing phiC31 integrase. Nat Biotechnol 21, 321–324. 10.1038/nbt787.

Bhatia, S., Kleinjan, D.J., Uttley, K., Mann, A., Dellepiane, N., and Bickmore, W.A. (2021). Quantitative spatial and temporal assessment of regulatory element activity in zebrafish. eLife 10. 10.7554/eLife.65601.

Bischof, J., Maeda, R.K., Hediger, M., Karch, F., and Basler, K. (2007). An optimized transgenesis system for Drosophila using germ-line-specific phiC31 integrases. Proc Natl Acad Sci U S A 104, 3312–3317. 10.1073/pnas.0611511104.

Boel, A., De Saffel, H., Steyaert, W., Callewaert, B., De Paepe, A., Coucke, P.J., and Willaert, A. (2018). CRISPR/Cas9-mediated homology-directed repair by ssODNs in zebrafish induces complex mutational patterns resulting from genomic integration of repair-template fragments. Dis Model Mech 11. 10.1242/dmm.035352.

Calos, M.P. (2006). The phiC31 integrase system for gene therapy. Curr Gene Ther 6, 633–645. 10.2174/156652306779010642.

Cui, X., Ji, D., Fisher, D.A., Wu, Y., Briner, D.M., and Weinstein, E.J. (2011). Targeted integration in rat and mouse embryos with zinc-finger nucleases. Nat Biotechnol 29, 64–67. nbt.1731 [pii] 10.1038/nbt.1731.

Cully, M. (2019). Zebrafish earn their drug discovery stripes. Nature reviews. Drug discovery 18, 811–813. 10.1038/d41573-019-00165-x.

Demarest, S.T., and Brooks-Kayal, A. (2018). From molecules to medicines: the dawn of targeted therapies for genetic epilepsies. Nat Rev Neurol 14, 735–745. 10.1038/s41582-018-0099-3.

DiNapoli, S.E., Martinez-McFaline, R., Gribbin, C.K., Wrighton, P.J., Balgobin, C.A., Nelson, I., Leonard, A., Maskin, C.R., Shwartz, A., Quenzer, E.D., et al. (2020). Synthetic CRISPR/Cas9 reagents facilitate genome editing and homology directed repair. Nucleic Acids Res 48, e38. 10.1093/nar/gkaa085.

Durrant, M.G.F., Alison; Tycko, Josh; Hinks, Michaela; Chandrasekaran, Sita S.; Perry, Nicholas T.; Schaepe, Julia; Du, Peter P.; Lotfy, Peter; Bassik, Michael C.; Bintu, Lacramioara; Bhatt2, Ami S.; Hsu, Patrick D. (2021). Large-scale discovery of recombinases for integrating DNA into the human genome. BioRxiv. https://doi.org/10.1101/2021.11.05.467528

Edwards, S.L., Beesley, J., French, J.D., and Dunning, A.M. (2013). Beyond GWASs: illuminating the dark road from association to function. Am J Hum Genet 93, 779–797. 10.1016/j.ajhg.2013.10.012.

Fazio, M., Ablain, J., Chuan, Y., Langenau, D.M., and Zon, L.I. (2020). Zebrafish patient avatars in cancer biology and precision cancer therapy. Nat Rev Cancer 20, 263–273. 10.1038/s41568-020-0252-3.

Foley, J.E., Maeder, M.L., Pearlberg, J., Joung, J.K., Peterson, R.T., and Yeh, J.R. (2009). Targeted mutagenesis in zebrafish using customized zinc-finger nucleases. Nat Protoc 4, 1855–1867. nprot.2009.209 [pii] 10.1038/nprot.2009.209.

Gagnon, J.A., Valen, E., Thyme, S.B., Huang, P., Akhmetova, L., Pauli, A., Montague, T.G., Zimmerman, S., Richter, C., and Schier, A.F. (2014). Efficient mutagenesis by Cas9 protein-mediated oligonucleotide insertion and large-scale assessment of single-guide RNAs. PLoS One 9, e98186. 10.1371/journal.pone.0098186.

Goldsmith, J.R., and Jobin, C. (2012). Think small: zebrafish as a model system of human pathology. J Biomed Biotechnol 2012, 817341. 10.1155/2012/817341.

Gusev, A., Mancuso, N., Won, H., Kousi, M., Finucane, H.K., Reshef, Y., Song, L., Safi, A., Schizophrenia Working Group of the Psychiatric Genomics, C., McCarroll, S., et al. (2018). Transcriptome-wide association study of schizophrenia and chromatin activity yields mechanistic disease insights. Nat Genet 50, 538–548. 10.1038/s41588-018-0092-1.

Gut, P., Reischauer, S., Stainier, D.Y.R., and Arnaout, R. (2017). Little Fish, Big Data: Zebrafish as a Model for Cardiovascular and Metabolic Disease. Physiol Rev 97, 889–938. 10.1152/physrev.00038.2016.

Helenius, I.T., and Yeh, J.R. (2012). Small zebrafish in a big chemical pond. J Cell Biochem 113, 2208–2216. 10.1002/jcb.24120.

Hillman, R.T., and Calos, M.P. (2012). Site-specific integration with bacteriophage PhiC31 integrase. Cold Spring Harb Protoc 2012. 10.1101/pdb.prot069211.

Hisano, Y., Sakuma, T., Nakade, S., Ohga, R., Ota, S., Okamoto, H., Yamamoto, T., and Kawahara, A. (2015). Precise in-frame integration of exogenous DNA mediated by CRISPR/Cas9 system in zebrafish. Sci Rep 5, 8841. 10.1038/srep08841.

Howe, K., Clark, M.D., Torroja, C.F., Torrance, J., Berthelot, C., Muffato, M., Collins, J.E., Humphray, S., McLaren, K., Matthews, L., et al. (2013). The zebrafish reference genome sequence and its relationship to the human genome. Nature 496, 498–503. 10.1038/nature12111.

Jao, L.E., Wente, S.R., and Chen, W. (2013). Efficient multiplex biallelic zebrafish genome editing using a CRISPR nuclease system. Proc Natl Acad Sci U S A 110, 13904–13909. 10.1073/pnas.1308335110.

Jusiak, B., Jagtap, K., Gaidukov, L., Duportet, X., Bandara, K., Chu, J., Zhang, L., Weiss, R., and Lu, T.K. (2019). Comparison of Integrases Identifies Bxb1-GA Mutant as the Most Efficient Site-Specific Integrase System in Mammalian Cells. ACS Synth Biol 8, 16–24. 10.1021/acssynbio.8b00089.

Kirchmaier, S., Hockendorf, B., Moller, E.K., Bornhorst, D., Spitz, F., and Wittbrodt, J. (2013). Efficient site-specific transgenesis and enhancer activity tests in medaka using PhiC31 integrase. Development 140, 4287–4295. 10.1242/dev.096081.

Li, Y.E., Allen, B.G., and Weeks, D.L. (2012). Using PhiC31 integrase to mediate insertion of DNA in Xenopus embryos. Methods Mol Biol 917, 219–230. 10.1007/978-1-61779-992-1_13.

Low, B.E., Hosur, V., Lesbirel, S., and Wiles, M.V. (2022). Efficient targeted transgenesis of large donor DNA into multiple mouse genetic backgrounds using bacteriophage Bxb1 integrase. Sci Rep 12, 5424. 10.1038/s41598-022-09445-w.

MacRae, C.A., and Peterson, R.T. (2015). Zebrafish as tools for drug discovery. Nature reviews. Drug discovery 14, 721–731. 10.1038/nrd4627.

Meyer, M., de Angelis, M.H., Wurst, W., and Kuhn, R. (2010). Gene targeting by homologous recombination in mouse zygotes mediated by zinc-finger nucleases. Proc Natl Acad Sci U S A 107, 15022–15026. 10.1073/pnas.1009424107.

Moreno-Mateos, M.A., Fernandez, J.P., Rouet, R., Vejnar, C.E., Lane, M.A., Mis, E., Khokha, M.K., Doudna, J.A., and Giraldez, A.J. (2017). CRISPR-Cpf1 mediates efficient homology-directed repair and temperature-controlled genome editing. Nat Commun 8, 2024. 10.1038/s41467-017-01836-2.

Mosimann, C., Kaufman, C.K., Li, P., Pugach, E.K., Tamplin, O.J., and Zon, L.I. (2011). Ubiquitous transgene expression and Cre-based recombination driven by the ubiquitin promoter in zebrafish. Development 138, 169–177. 10.1242/dev.059345.

Mosimann, C., Puller, A.C., Lawson, K.L., Tschopp, P., Amsterdam, A., and Zon, L.I. (2013a). Site-directed zebrafish transgenesis into single landing sites with the phiC31 integrase system. Developmental dynamics: an official publication of the American Association of Anatomists 242, 949–963. 10.1002/dvdy.23989.

Mosimann, C., Puller, A.C., Lawson, K.L., Tschopp, P., Amsterdam, A., and Zon, L.I. (2013b). Site-directed zebrafish transgenesis into single landing sites with the phiC31 integrase system. Developmental dynamics: an official publication of the American Association of Anatomists 242, 949–963. 10.1002/dvdy.23989.

Petri, K., Zhang, W., Ma, J., Schmidts, A., Lee, H., Horng, J.E., Kim, D.Y., Kurt, I.C., Clement, K., Hsu, J.Y., et al. (2022). CRISPR prime editing with ribonucleoprotein complexes in zebrafish and primary human cells. Nat Biotechnol 40, 189–193. 10.1038/s41587-021-00901-y.

Prill, K., and Dawson, J.F. (2020). Homology-Directed Repair in Zebrafish: Witchcraft and Wizardry? Front Mol Biosci 7, 595474. 10.3389/fmolb.2020.595474.

Prykhozhij, S.V., Fuller, C., Steele, S.L., Veinotte, C.J., Razaghi, B., Robitaille, J.M., McMaster, C.R., Shlien, A., Malkin, D., and Berman, J.N. (2018a). Optimized knock-in of point mutations in zebrafish using CRISPR/Cas9. Nucleic Acids Res 46, 9252. 10.1093/nar/gky674.

Prykhozhij, S.V., Fuller, C., Steele, S.L., Veinotte, C.J., Razaghi, B., Robitaille, J.M., McMaster, C.R., Shlien, A., Malkin, D., and Berman, J.N. (2018b). Optimized knock-in of point mutations in zebrafish using CRISPR/Cas9. Nucleic Acids Res 46, e102. 10.1093/nar/gky512.

Raymond, C.S., and Soriano, P. (2007). High-efficiency FLP and PhiC31 site-specific recombination in mammalian cells. PLoS One 2, e162. 10.1371/journal.pone.0000162.

Richardson, C.D., Ray, G.J., DeWitt, M.A., Curie, G.L., and Corn, J.E. (2016). Enhancing homology-directed genome editing by catalytically active and inactive CRISPR-Cas9 using asymmetric donor DNA. Nat Biotechnol 34, 339–344. 10.1038/nbt.3481.

Rickert, R.C., Roes, J., and Rajewsky, K. (1997). B lymphocyte-specific, Cre-mediated mutagenesis in mice. Nucleic Acids Res 25, 1317–1318. 10.1093/nar/25.6.1317.

Rissone, A., and Burgess, S.M. (2018). Rare Genetic Blood Disease Modeling in Zebrafish. Front Genet 9, 348. 10.3389/fgene.2018.00348.

Roberts, J.A., Miguel-Escalada, I., Slovik, K.J., Walsh, K.T., Hadzhiev, Y., Sanges, R., Stupka, E., Marsh, E.K., Balciuniene, J., Balciunas, D., and Muller, F. (2014). Targeted transgene integration overcomes variability of position effects in zebrafish. Development 141, 715–724. 10.1242/dev.100347.

Sakai, C., Ijaz, S., and Hoffman, E.J. (2018). Zebrafish Models of Neurodevelopmental Disorders: Past, Present, and Future. Front Mol Neurosci 11, 294. 10.3389/fnmol.2018.00294.

Shin, J., Chen, J., and Solnica-Krezel, L. (2014). Efficient homologous recombination-mediated genome engineering in zebrafish using TALE nucleases. Development 141, 3807–3818. 10.1242/dev.108019.

Soriano, P. (1999). Generalized lacZ expression with the ROSA26 Cre reporter strain. Nat Genet 21, 70–71. 10.1038/5007.

Swinney, D.C., and Anthony, J. (2011). How were new medicines discovered? Nature reviews. Drug discovery 10, 507–519. 10.1038/nrd3480.

Thorpe, H.M., and Smith, M.C. (1998). In vitro site-specific integration of bacteriophage DNA catalyzed by a recombinase of the resolvase/invertase family. Proc Natl Acad Sci U S A 95, 5505–5510. 10.1073/pnas.95.10.5505.

Torraca, V., and Mostowy, S. (2018). Zebrafish Infection: From Pathogenesis to Cell Biology. Trends Cell Biol 28, 143–156. 10.1016/j.tcb.2017.10.002.

Traver, D., Paw, B.H., Poss, K.D., Penberthy, W.T., Lin, S., and Zon, L.I. (2003). Transplantation and in vivo imaging of multilineage engraftment in zebrafish bloodless mutants. Nat Immunol 4, 1238–1246. 10.1038/ni1007.

Tucker, N.R., Dolmatova, E.V., Lin, H., Cooper, R.R., Ye, J., Hucker, W.J., Jameson, H.S., Parsons, V.A., Weng, L.C., Mills, R.W., et al. (2017). Diminished PRRX1 Expression Is Associated With Increased Risk of Atrial Fibrillation and Shortening of the Cardiac Action Potential. Circ Cardiovasc Genet 10. 10.1161/CIRCGENETICS.117.001902.

Vacaru, A.M., Unlu, G., Spitzner, M., Mione, M., Knapik, E.W., and Sadler, K.C. (2014). In vivo cell biology in zebrafish -providing insights into vertebrate development and disease. J Cell Sci 127, 485–495. 10.1242/jcs.140194.

Wierson, W.A., Welker, J.M., Almeida, M.P., Mann, C.M., Webster, D.A., Torrie, M.E., Weiss, T.J., Kambakam, S., Vollbrecht, M.K., Lan, M., et al. (2020). Efficient targeted integration directed by short homology in zebrafish and mammalian cells. eLife 9. 10.7554/eLife.53968.

Yang, H., Ren, S., Yu, S., Pan, H., Li, T., Ge, S., Zhang, J., and Xia, N. (2020). Methods Favoring Homology-Directed Repair Choice in Response to CRISPR/Cas9 Induced-Double Strand Breaks. Int J Mol Sci 21. 10.3390/ijms21186461.

